# An ancient thyrostimulin-like neuroendocrine pathway regulates growth in *Caenorhabditis elegans*

**DOI:** 10.1101/2023.02.16.528858

**Authors:** Signe Kenis, Majdulin Nabil Istiban, Sara Van Damme, Elke Vandewyer, Jan Watteyne, Liliane Schoofs, Isabel Beets

## Abstract

In vertebrates, thyrostimulin is a highly conserved glycoprotein hormone that, besides thyroid stimulating hormone (TSH), is a potent ligand of the TSH receptor. Thyrostimulin is considered the most ancestral glycoprotein hormone and orthologs of its subunits, GPA2 and GPB5, are widely conserved across vertebrate and invertebrate animals. Unlike TSH, however, the functions of the thyrostimulin neuroendocrine system remain largely unexplored. Here, we identify a functional thyrostimulin-like signaling system in *Caenorhabditis elegans*. We show that orthologs of GPA2 and GPB5, together with thyrotropin-releasing hormone (TRH) related neuropeptides, constitute a neuroendocrine pathway that promotes growth in *C. elegans*. GPA2/GPB5 signaling is required for normal body size and acts through activation of the glycoprotein hormone receptor ortholog FSHR-1. *C. elegans* GPA2 and GPB5 increase cAMP signaling by FSHR-1 *in vitro*. Both subunits are expressed in enteric neurons and promote growth by signaling to their receptor in glial cells and the intestine. Impaired GPA2/GPB5 signaling causes bloating of the intestinal lumen. In addition, mutants lacking thyrostimulin-like signaling are defective in defecation behavior. Our study suggests that the thyrostimulin GPA2/GPB5 pathway is an ancient enteric neuroendocrine system that regulates intestinal function in ecdysozoans, and may ancestrally have been involved in the control of organismal growth.

## Introduction

Glycoprotein hormones are key neuroendocrine factors that control diverse physiological processes, such as development, reproduction, energy homeostasis and growth ^1,2^. In vertebrates, five glycoprotein hormones have been described that are all formed by the heterodimerization of two cysteine-knot containing polypeptides, an alpha (GPA) and beta (GPB) subunit, which are non-covalently associated ^1,3–5^. Four of them, including thyroid-stimulating hormone (TSH), follicle-stimulating hormone (FSH), luteinizing hormone (LH) and chorionic gonadotropin (CG), share a common alpha subunit, GPA1, that associates with hormone-specific beta subunits (TSHβ, FSHβ, LHβ and CGβ) ^3,6,7^. A fifth glycoprotein hormone, called thyrostimulin, was discovered more recently and is comprised of a unique glycoprotein alpha 2 (GPA2) and glycoprotein beta 5 (GPB5) subunit ^8,9^. Thyrostimulin GPA2 and GPB5 subunits are widely conserved across bilaterian animals, whereas the conservation of GPA1 and other GPB polypeptides is restricted to vertebrates ^1,8,10^. Orthologs of GPA2 and GPB5 have been identified both in vertebrates and invertebrates, including arthropods ^11–14^, nematodes ^9,15^, annelids ^16,17^,mollusks ^18–20^, urochordates ^21,22^, and cephalochordates ^21,23,24^. Thyrostimulin is therefore considered the most ancestral glycoprotein hormone, from which other vertebrate glycoprotein hormone systems most likely evolved through gene duplication ^3,7,8^.

In contrast to the well-studied TSH and gonadotropin endocrine systems, the physiological role of thyrostimulin remains largely unexplored. In 2002, a receptor for human thyrostimulin was identified ^10^. The heterodimeric GPA2/GPB5 glycoprotein has been shown to activate the human TSH receptor (TSHR), but not FSH or LH/CG receptors, and is capable of inducing receptor-mediated cyclic adenosine monophosphate (cAMP) production more potently than human TSH itself ^10,25^. Like TSH, GPA2/GPB5 increases thyroid hormone secretion in rats and, based on its TSHR- and thyroid-stimulating activity, was named thyrostimulin ^10^. Later studies in basal vertebrates, urochordates, cephalochordates, and insects revealed that the interaction of thyrostimulin with TSHR orthologs is widely conserved ^14,23,26– 29^. Glycoprotein hormone receptors belong to a subfamily of leucine-rich repeat containing G protein-coupled receptors (LGRs) that are conserved across divergent animal phyla ^7,8,30,31^. Thyrostimulin orthologs were found to activate LGRs related to TSHRs in lampreys, tunicates, *Amphioxus*, the fruit fly *Drosophila melanogaster* and the mosquito *Aedes aegypti* ^14,23,26–29^, indicating that the evolutionary origin of this glycoprotein hormone-receptor system dates back to a common ancestor of bilaterian animals.

To date, few studies have investigated the physiological role of thyrostimulin in vertebrates. These suggest that thyrostimulin has pleiotropic functions, as it is involved in reproduction ^32^, skeletal development ^33,34^, immune responses ^35,36^ and thyroxine production ^10,37^. In invertebrates, the functions of thyrostimulin-like signaling have so far been studied only in arthropods. GPA2/GPB5 or its receptor have been implicated in the control of reproduction, development, feeding, diuresis and osmoregulation in insects ^12,38–40^. In the prawn *Macrobrachium rosenberghii*, knockdown of GPA2/GPB5 or its predicted receptor affects female reproduction ^41^. The mechanisms by which GPA2/GPB5 regulates these biological processes as well as its suggested hormonal functions in other invertebrates, however, are still undetermined.

Due to its well-defined nervous system and extensive genetic toolbox, the nematode *Caenorhabditis elegans* is an excellent model to study neuroendocrine systems. Several vertebrate endocrine factors, like gonadotropin-releasing hormone (GnRH) and thyrotropin-releasing hormone (TRH), have been identified in *C. elegans* and were found to have conserved physiological roles, such as in the control of reproduction and growth ^42,43^. Phylogenetic analyses also revealed orthologs of the thyrostimulin GPA2 and GPB5 subunits in the *C. elegans* genome ^9,15,21,40^, which carry a cysteine knot motif of six cysteine residues that is typical of glycoprotein hormone subunits ^44,45^. Recent analysis of the crystal structure of *C. elegans* GPA2/GPB5 validates its structural similarity to vertebrate glycoprotein hormones that was previously found by comparative genomic analyses ^3,8,46^. In addition, *C. elegans* has only one receptor ortholog of the vertebrate glycoprotein hormone receptors. This *C. elegans* receptor, designated as FSHR-1, belongs to the same LGR subfamily as the vertebrate thyrostimulin receptors ^7,8^, but is still orphan. Here, we identify *C. elegans* GPA2 and GPB5 as ligands of FSHR-1. Using CRISPR/Cas9 reverse genetics, we functionally characterize the thyrostimulin-related system in *C. elegans* and demonstrate that GPA2/GPB5 signaling, together with TRH-related neuropeptides, regulates growth and intestinal function.

## Results

### *C. elegans* thyrostimulin GPA2/GPB5 orthologs regulate growth

Phylogenetic analysis revealed two orthologs of thyrostimulin GPA2 and GPB5 subunits in the *C. elegans* genome ^8,9^. These are encoded by the genes *flr-2* and *T23B12*.*8* ^15,21^, which we called *gpla-1* and *gplb-1* (glycoprotein hormone-like alpha and beta), respectively. GPLA-1 and GPLB-1 resemble vertebrate GPA2 and GPB5 and are widely conserved in nematodes (Fig. S1 and S2). Although the roles of thyrostimulin are not well understood, GPA2/GPB5 signaling in vertebrates and insects regulates several aspects of physiology, including growth, development and intestinal functioning ^11,13,15,18,38,40,47^. In *C. elegans*, mutations in *gpla-1* have been reported to suppress slow growing of several growth-defective mutants ^15,48^. We therefore hypothesized that thyrostimulin-like signaling may influence *C. elegans* body size. To test this, we generated complete gene knockouts of *gpla-1* and *gplb-1* using CRISPR/Cas9 gene editing (Fig. 1A) and investigated if these null mutants show body size defects at day 1 of adulthood (65 hours post L1 arrest). The generated *gpla-1 (ibt1)* and *gplb-1 (ibt4)* mutants display a significantly shorter body length compared to wild-type animals (Fig. 1B-C). This size defect persists for at least 5 days into adulthood, suggesting that it results from defective growth rather than a delay in the timing of development (Fig. S3A). Additionally, worms lacking *gpla-1* and *gplb-1* have a reduced body width compared to wild type (Fig. S3B). We also measured body sizes of two independently generated *gpla-1* mutants, carrying the *ut5* and *ok2127* alleles, and found that they also display a decreased body length compared to wild-type animals. Mutants carrying the *ut5* allele show a reduction in body width as well (Fig. S3C-D). The defect in length, however, is most pronounced in all *gpla-1* and *gplb-1* mutants. Therefore, we quantified body length in further experiments. Expressing wild-type *gpla-1* and *gplb-1* in their respective mutant background fully restored body length (Fig. 1B-C), indicating that both GPA2 and GPB5 orthologs are required for normal body size.

**Figure 1.**
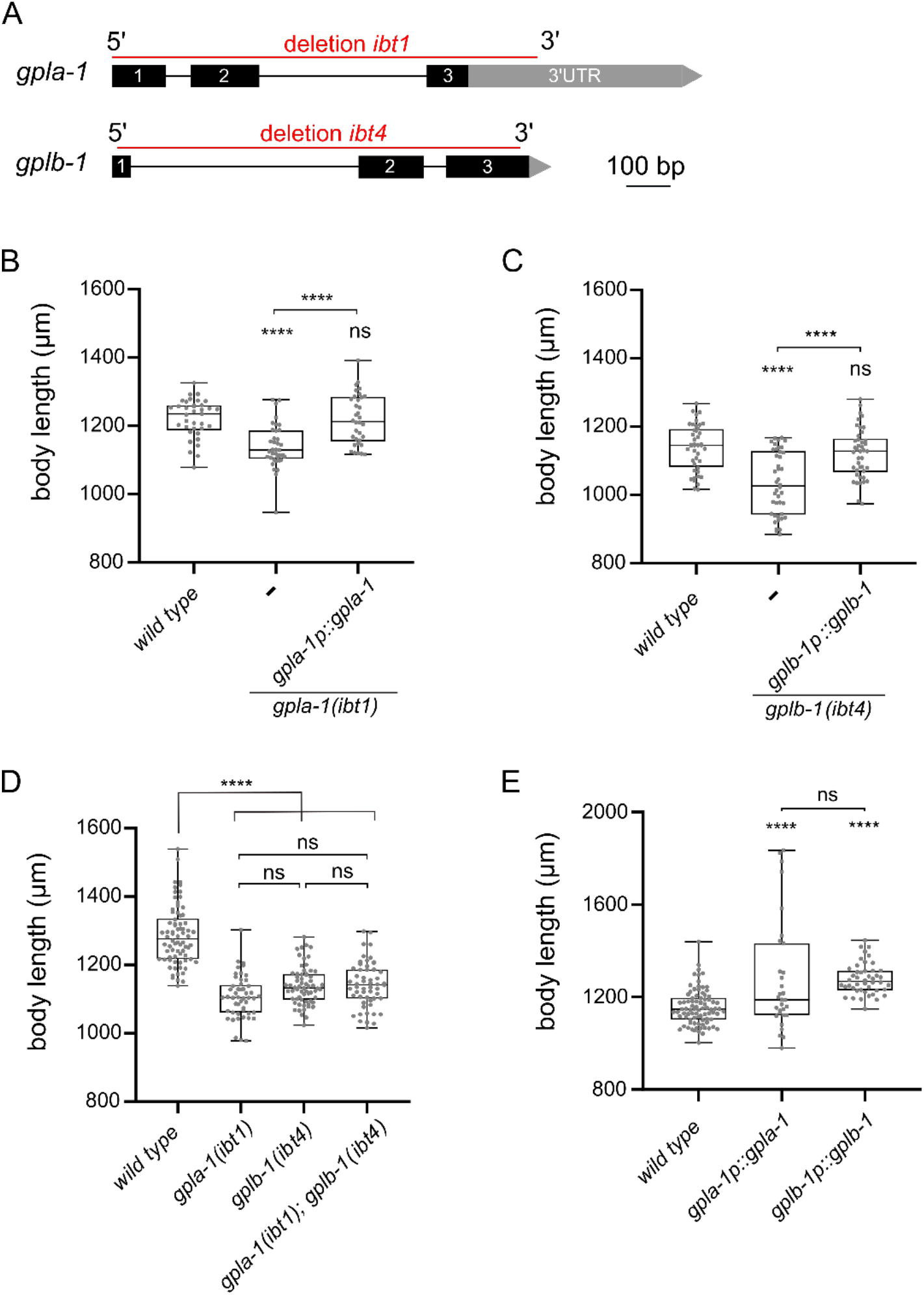
Thyrostimulin-like signaling promotes *C. elegans* growth. (A) Structure of *gpla-1* and *gplb-1* transcripts. Black boxes represent exons; black lines indicate intronic regions; and grey boxes and arrows represent 3’UTR sequences. Red lines mark the positions of the *ibt1* and *ibt4* deletions created by CRISPR/Cas9 gene editing. (B-C) Null mutants of *gpla-1* and *gplb-1* have a significantly shorter body length than wild-type animals at day 1 of adulthood. Restoring wild-type *gpla-1* (B) and *gplb-1* (C) expression rescues the body size defect. (D) Double mutants of *gpla-1* and *gplb-1* do not significantly differ in body length from the single mutants. (E) Overexpressing wild-type *gpla-1* and *gplb-1* significantly increases body length compared to wild type. Boxplots indicate 25^th^ (lower boundary), 50^th^ (central line), and 75^th^ (upper boundary) percentiles. Whiskers show minimum and maximum values. For (B) – (E), each genotype was tested in at least 2 assays with 10–30 animals per trial. Data were analyzed by one-way ANOVA with Tukey’s multiple comparisons test. ****P < 0.0001; ns, not significant.

To determine whether GPLA-1 and GPLB-1 act in the same growth-regulating pathway, we quantified the body length of double mutants. Mutants lacking both genes did not significantly differ in length from the single mutants (Fig. 1D). This suggests that *gpla-1* and *gplb-1* indeed act in the same genetic pathway to control body size. Overexpression of each individual gene increased the length of wild-type worms (Fig. 1E), suggesting that GPLA-1 and GPLB-1 are sufficient to induce growth.

### GPLA-1/GPA2 and GPLB-1/GPB5 control body size via the glycoprotein hormone receptor ortholog FSHR-1

To unravel how thyrostimulin-like signaling regulates *C. elegans* body size, we aimed to identify its target receptor. Since glycoprotein hormones typically bind to LGRs ^8^, we hypothesized that GPLA-1 and GPLB-1 may signal via FSHR-1, which is the sole glycoprotein hormone receptor ortholog in *C. elegans* ^49^. To determine whether *C. elegans* thyrostimulin can activate FSHR-1, we used a cAMP-based receptor activation assay in human embryonic kidney (HEK) cells. We co-expressed FSHR-1 with a cAMP response element (CRE)-luciferase reporter in HEK cells and quantified its activation by assessing bioluminescence levels in the presence of luciferin substrate (Fig. 2A). Consistent with previous reports ^49^, we observed basal activity of FSHR-1, as bioluminescent cAMP signals were higher in receptor-expressing cells than in cells transfected with an empty control vector (Fig. S4A). However, challenging cells with recombinant GPLA-1 and GPLB-1 proteins significantly increased cAMP signaling by FSHR-1 (Fig. 2B-D). To generate recombinant GPLA-1/GPLB-1, HEK293 cells were first transfected with expression vectors encoding His-tagged GPLA-1 and non-tagged GPLB-1. Tagging of GPLA-1 did not affect protein function as transgenic *C. elegans* expressing endogenous His-GPLA-1 showed normal body size (Fig. S4B). Recombinant proteins purified from cells expressing both subunits significantly increased cAMP signaling by FSHR-1 (Fig. 2B). Although we could not confirm the heterodimeric nature of recombinant proteins due to the limited yield, this suggests that GPLA-1/GPLB-1 is a ligand of FSHR-1. We also tested whether His-tagged GPLA-1 and GPLB-1 proteins could activate FSHR-1 individually. For this, recombinant proteins were purified from HEK293 cells transfected with an expression vector encoding only one of the two subunits. We found that GPLA-1 as well as GPLB-1 increased cAMP signaling by FSHR-1 (Fig. 2C and D), which indicates that each subunit is capable of activating the receptor. None of the purified recombinant proteins elicited a significant cAMP response in cells transfected with an empty control vector instead of the FSHR-1 expression plasmid (Fig. S4C). Taken together, these results suggest that GPLA-1 and GPLB-1 are cognate ligands of FSHR-1.

**Figure 2.**
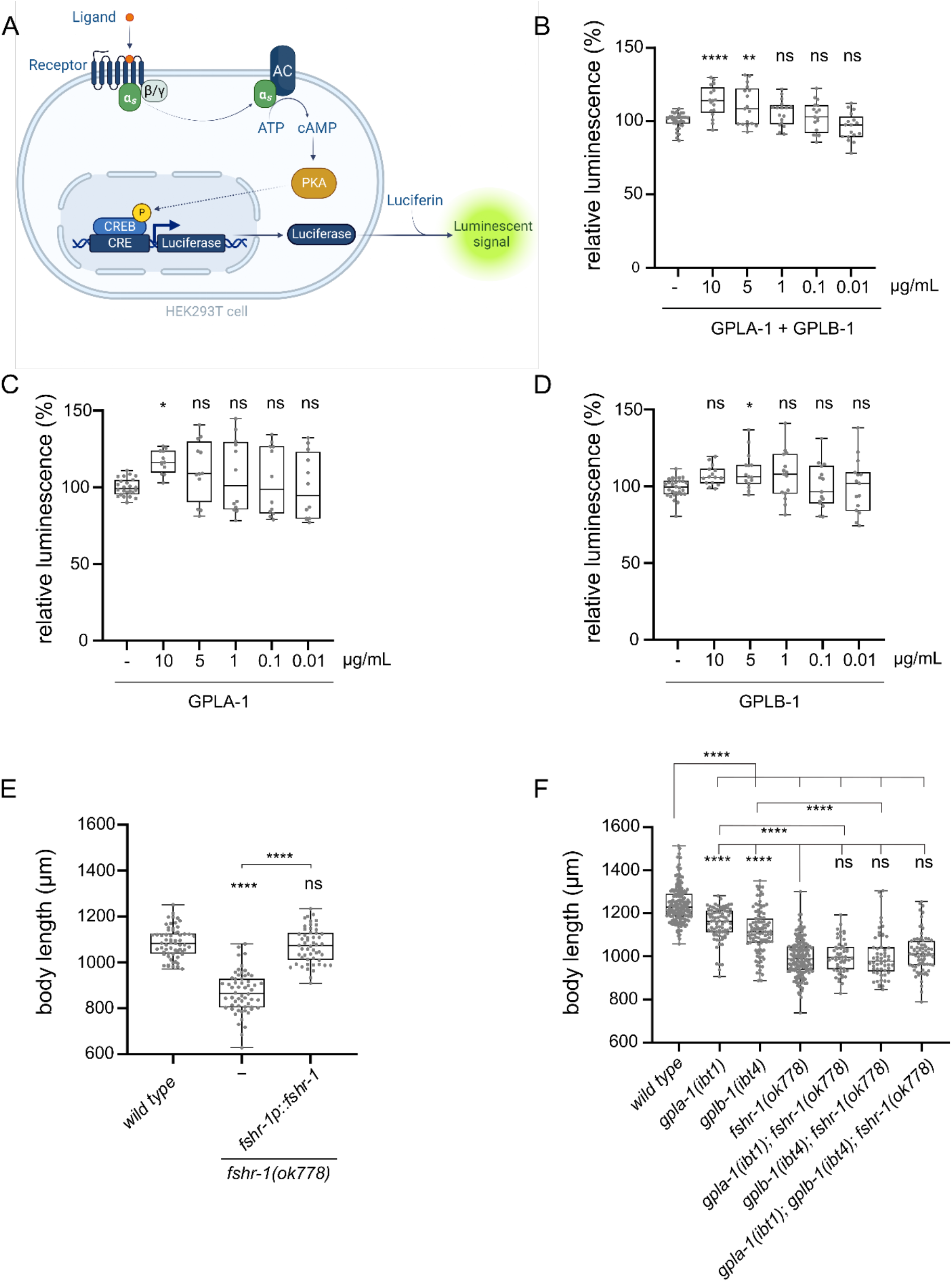
*C. elegans* GPLA-1 and GPLB-1 regulate growth through the glycoprotein hormone receptor ortholog FSHR-1. (A) Luminescence-based cAMP assay for measuring GPCR activation. FSHR-1 is co-expressed with the CRE(6x)-luciferase reporter in HEK cells. Receptor activation initiates the production of cAMP by adenylyl cyclase (AC), which is monitored by the cAMP-sensitive biosensor luciferase in the presence of luciferin substrate. Created with BioRender.com. (B-D) Recombinant GPLA-1 and GPLB-1 proteins activate FSHR-1 in a cellular assay described in panel A. Data are shown as relative luminescence after normalization to the ligand-free control. One-way ANOVA with Dunnett’s multiple comparisons test. n ≥ 4 assays. (E) *fshr-1* mutants are significantly shorter than wild-type animals, which is rescued by expressing wild-type *fshr-1* under its endogenous promoter. (F) Double and triple mutants of *gpla-1, gplb-1* and *fshr-1* do not significantly differ in body length from *fshr-1* single mutants. Body size data were analyzed by one-way ANOVA with Tukey’s multiple comparisons test. Boxplots indicate 25^th^ (lower boundary), 50^th^ (central line), and 75^th^ (upper boundary) percentiles. Whiskers show minimum and maximum values. For (E) and (F), each genotype was tested in at least 3 assays with 20-30 animals per trial. *P < 0.05; **P < 0.01; ****P < 0.0001; ns, not significant.

As FSHR-1 is activated by GPLA-1 and GPLB-1 *in vitro*, we asked if this receptor is involved in growth regulation. We observed that a loss-of-function mutant of *fshr-1* suffers from severe growth defects that persist throughout adulthood (Fig. 2E and S3A). Expressing wild-type *fshr-1* fully restores this growth defect, indicating that this receptor, like its activating ligands GPLA-1 and GPLB-1, is required for growth (Fig. 2E). Although body length was more strongly reduced in *fshr-1* animals than in mutants lacking the thyrostimulin subunits, double and triple mutants of *gpla-1, gplb-1* and their receptor *fshr-1* were not shorter than the *fshr-1* single mutant (Fig. 2F). This suggests they all act in the same growth-regulating pathway *in vivo*, consistent with our finding that FSHR-1 is a receptor for GPLA-1 and GPLB-1 *in vitro*.

### *C. elegans* GPA2/GPB5 and FSHR-1 are expressed in enteric neurons and the intestine

To gain further insight into the role of thyrostimulin-like signaling in growth regulation, we investigated the expression patterns of *gpla-1, gplb-1* and *fshr-1*. Fluorescent reporter transgenes, which rescue body length defects, showed expression in multiple pharyngeal neurons and tissues associated with feeding and digestive processes. Consistent with single-cell RNA sequencing data and previous reporter analyses ^15,50,51^, we found that *gpla-1* is predominantly expressed in neurons associated with the gastrointestinal tract (Fig. 3A-B), which are part of the enteric nervous system in *C. elegans* ^51–53^. These include the pharyngeal neurons M1, M5, I5 and NSM, as well as the excitatory motor neurons AVL and DVB that control enteric muscle contractions in the hindgut during defecation ^54–56^. We also observed expression of *gplb-1* in the enteric DVB neuron (Fig. 3D). In addition, both *gpla-1* and *gplb-1* reporter transgenes show fluorescence in the RME motor neurons that innervate the head muscles (Fig. 3A, C). Using a reporter transgene for GABAergic neurons, we confirmed that *gpla-1* and *gplb-1* are both expressed in the GABAergic DVB and RME neurons (Fig. S5). In addition, the *gplb-1* reporter transgene shows expression in non-neuronal tissues, including the head mesodermal cell (hmc) and the enteric muscles of the hindgut, which are involved in defecation ^54–58^.

**Figure 3.**
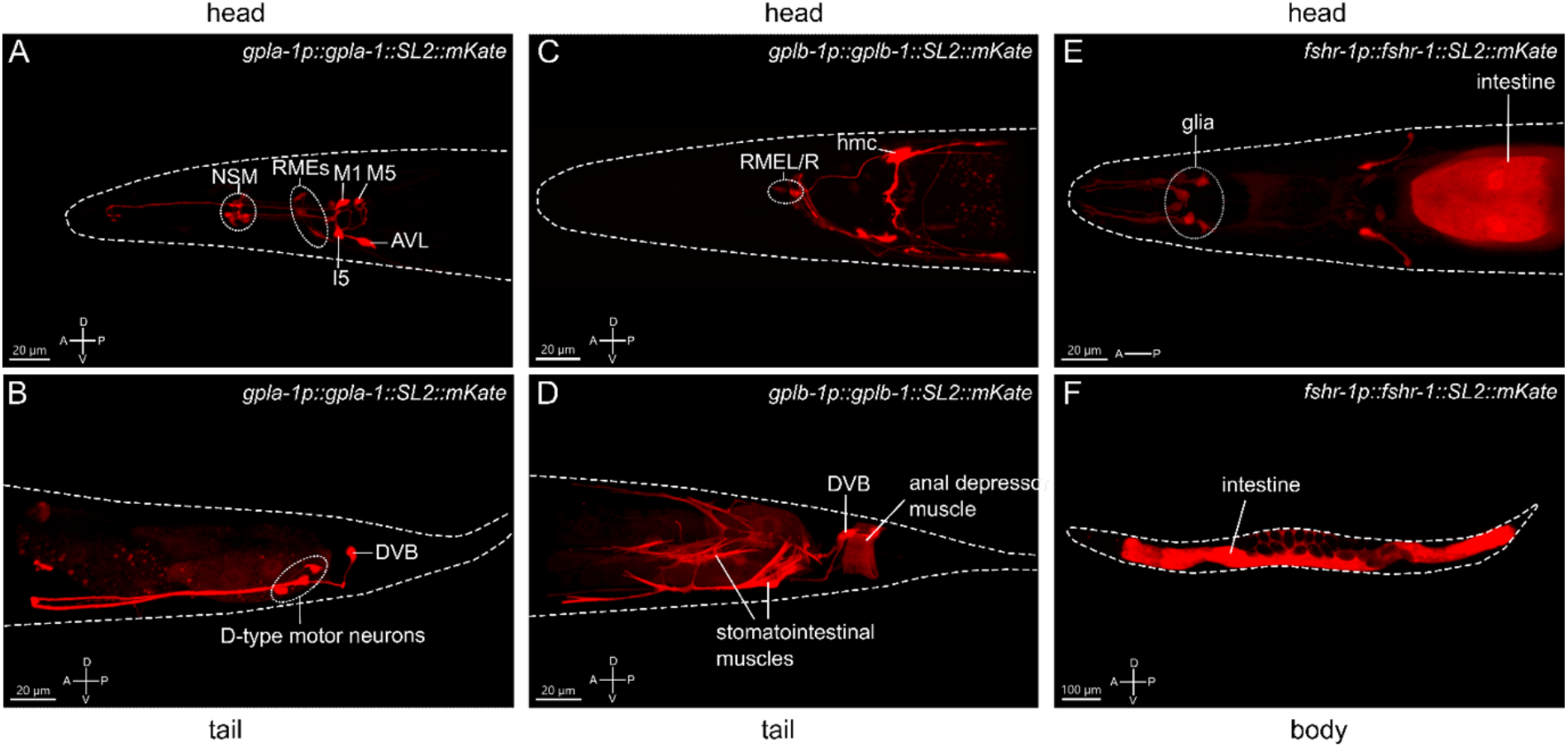
Expression patterns of the *C. elegans* thyrostimulin-like subunits GPLA-1/GPA2 and GPLB-1/GPB5 and their receptor FSHR-1. Representative confocal Z-stack projections of the regions showing expression of an mKate reporter transgene for *gpla-1* (A-B), *gplb-1* (C-D) and *fshr-1* (E-F). A; anterior, P; posterior, D; dorsal, V; ventral orientation.

A reporter construct for *fshr-1* recapitulated the reported expression of the receptor in the intestine and in multiple neurons in the head ^47,50,59,60^. In addition, we observed expression of *fshr-1* in several cells close to the anterior pharyngeal bulbus that, based on position and morphology of their projections, are most likely glial cells (Fig. 3E-F). Together, these expression patterns suggest that thyrostimulin-like signaling may influence body size by controlling intestinal or neural function.

### Intestinal and glial glycoprotein hormone receptor signaling controls body size

Next, we asked in which tissues *fshr-1* is required for normal body size. Expressing wild-type *fshr-1* under control of the intestine-specific *ges-1* ^61–63^ or the pan-glial *mir-228* ^64^ promoters fully rescued the body size defect of *fshr-1* mutants (Fig. 4A). By contrast, pan-neuronal expression using the *rab-3* ^65,66^ promoter partially restored body size (Fig. 4A). Since additional expression of this promoter has been reported in the intestine ^67,68^, we generated a second transgene for pan-neuronal expression of wild-type *fshr-1* using the *rgef-1* promoter ^69^. This transgene did not restore the body size defect of *fshr-1* mutants (Fig. 4A). Taken together, our results show that FSHR-1 signaling in the intestine and glial cells is required for normal body size.

**Figure 4.**
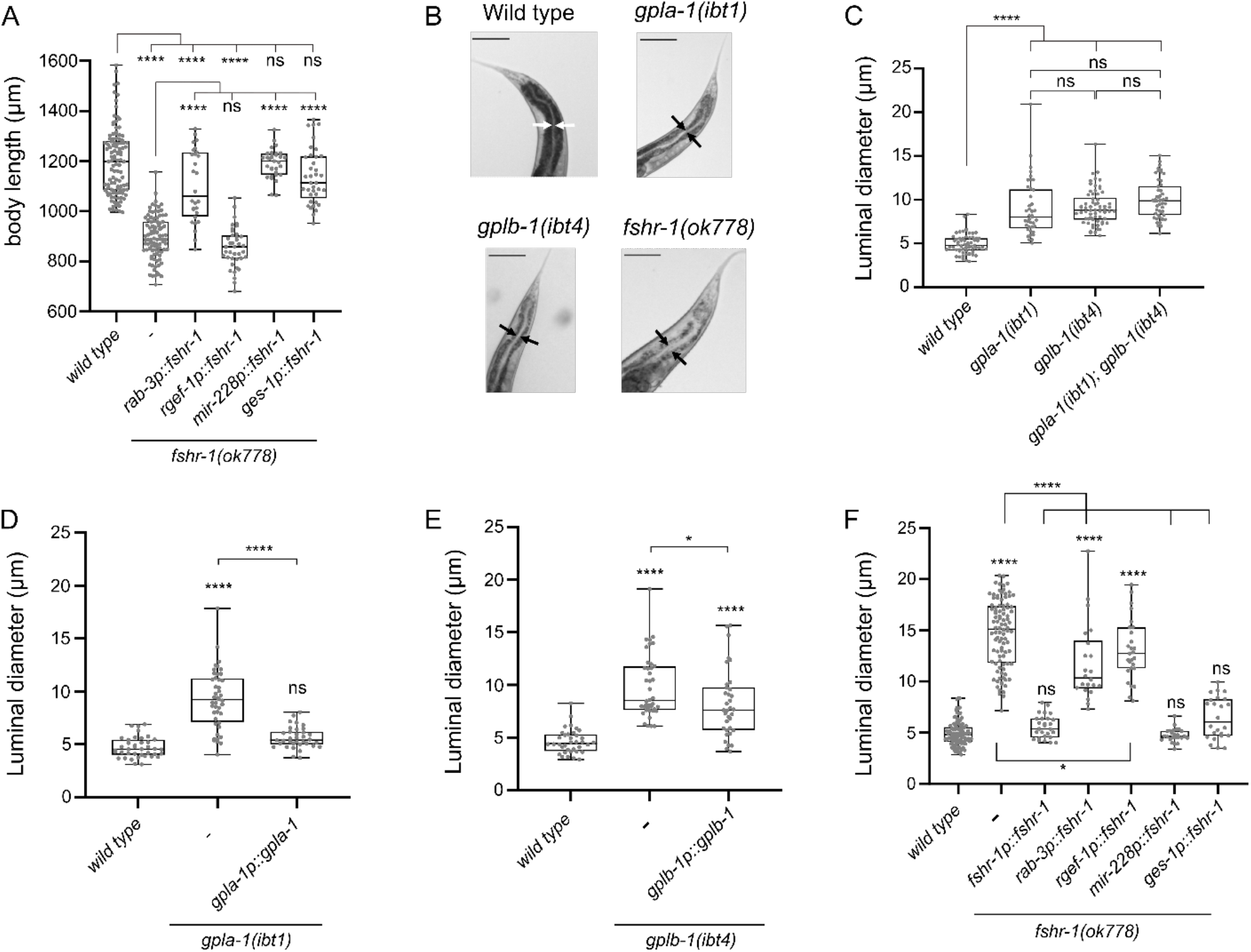
Thyrostimulin-like signaling promotes growth by regulating intestinal function. (A) Expression of wild-type *fshr-1* under control of the pan-glial promoter *mir-228p* or the intestinal promoter *ges-1p* fully restores the body size defect of *fshr-1* mutants. Expression of wild-type *fshr-1* under the pan-neuronal *rab-3* promoter partially rescues body size, whereas expression from the pan-neuronal *rgef-1* promoter does not. (B) Mutants deficient in thyrostimulin signaling show intestinal bloating. Scale bars represent 100 μm. (C) The intestinal lumen of *gpla-1* and *gplb-1* single and double mutants is significantly wider than that of wild-type animals. (D-E) Restoring wild-type *gpla-1* (D) and *gplb-1* (E) expression rescues the intestinal bloating phenotype. (F) Luminal size of the intestine is significantly increased in *fshr-1* mutants, and expressing wild-type copies of *fshr-1* under control of an endogenous promoter (*fshr-1p*), pan-glial promoter (*mir-228p*), or intestinal promoter (*ges-1p*) fully rescues intestinal bloating. Pan-neuronal *fshr-1* expression controlled by *rab-3p* or *rgef-1p* partially restores luminal size of the intestine. Boxplots indicate 25^th^ (lower boundary), 50^th^ (central line), and 75^th^ (upper boundary) percentiles. Whiskers show minimum and maximum values. For (A) and (C)-(F), each genotype was tested in at least 2 assays with 20-30 animals per trial. Data were analyzed by one-way ANOVA with Tukey’s multiple comparisons test. *P < 0.05; ****P < 0.0001; ns, not significant.

### Mutants deficient in thyrostimulin-like signaling show defecation defects and intestinal bloating

Our expression analysis and rescue experiments suggest that thyrostimulin-like signaling regulates intestinal function. To test this hypothesis, we investigated the anatomy of the intestine and measured the size of the intestinal lumen. The intestinal lumen was bloated to a similar extent in single and double mutants of *gpla-1* and *gplb-1* (Fig. 4B and C). The increased luminal size was fully rescued by expressing wild-type copies of *gpla-1* and partially restored by expressing wild-type *gplb-1* (Fig. 4D and E). Mutants lacking *fshr-1* also displayed a severely bloated intestinal lumen, which was rescued by restoring wild-type *fshr-1* expression under its endogenous promoter (Fig. 4B and F). Tissue-specific expression of wild-type *fshr-1* in the intestine or glial cells also fully rescued luminal bloating, whereas pan-neuronal expression only partially restored the luminal size (Fig. 4F). The lumen of double and triple mutants of *gpla-1, gplb-1* and their receptor *fshr-1* was not wider than that of the *fshr-1* single mutant (Fig. S6D-F), suggesting that they act in the same pathway regulating luminal bloating.

Since *gpla-1* and *gplb-1* are predominantly expressed in enteric neurons and tissues controlling the defecation motor program, we hypothesized that thyrostimulin signaling may influence growth by regulating defecation behavior. To test this, we first examined whether mutants defective in the defecation motor program also show growth defects. In *C. elegans*, the defecation motor program is a rhythmic behavior with a period of about 50 seconds, which ultimately results in contraction of the enteric muscles in the hindgut and the subsequent expulsion of digested food from the intestine ^54–56^. Mutants for *unc-25* and *exp-1* are deficient in synaptic GABAergic transmission, which drives enteric muscle contraction, and show severe defecation defects ^70–72^. We found that *unc-25(e156)* and *exp-1(ox276)* mutants are also significantly smaller than wild-type animals (Fig. S6A), suggesting that defecation defects correlate with impaired growth in these mutants. Next, we asked whether mutants defective in thyrostimulin-like signaling show defects in defecation behavior. Although the contraction frequency of the anterior body wall muscles (aBoc) did not differ from wild-type aBoc frequency (Fig. S6B), the average length of the defecation cycle was significantly longer in mutants for *gpla-1, gplb-1*, and *fshr-1* compared to the cycle length of wild-type animals (Fig. S6C). Taken together, these findings indicate that thyrostimulin-like neuroendocrine signaling regulates growth by controlling intestinal function.

### GPA2/GPB5 and TRH-like neuropeptide signaling act in the same pathway regulating growth

In vertebrates, glycoprotein hormone signaling is regulated by hypothalamic factors that control the release of hormones from the pituitary ^2,73^. Thyrotropin-releasing hormone (TRH) is a highly conserved releasing factor that regulates growth and metabolism across bilaterian animals ^2,43,73–75^. In *C. elegans* TRH-related neuropeptides promote growth and mutants lacking the TRH-like precursor gene *trh-1* have a smaller body size than wild-type animals ^43^. We therefore asked whether TRH- and thyrostimulin-like signaling systems may interact to control *C. elegans* growth.

To test this hypothesis, we compared the body size of single and double mutants lacking *trh-1* and the thyrostimulin subunit orthologs *gpla-1* and *gplb-1*. Double mutants defective in both signaling systems did not significantly differ in length compared to the single mutants (Fig. 5A). Similar to thyrostimulin mutants, worms lacking TRH-1 also displayed intestinal bloating (Fig. 5B and C). Although the effect is more severe in mutants lacking thyrostimulin-like signaling, the extent of luminal bloating in double mutants of *gpla-1, gplb-1* and *trh-1* did not significantly differ from single *gpla-1* or *gplb-1* mutants (Fig. 5D and E). These results indicate that *trh-1* and *gpla-1/gplb-1* act in the same growth-stimulating pathway *in vivo*.

**Figure 5.**
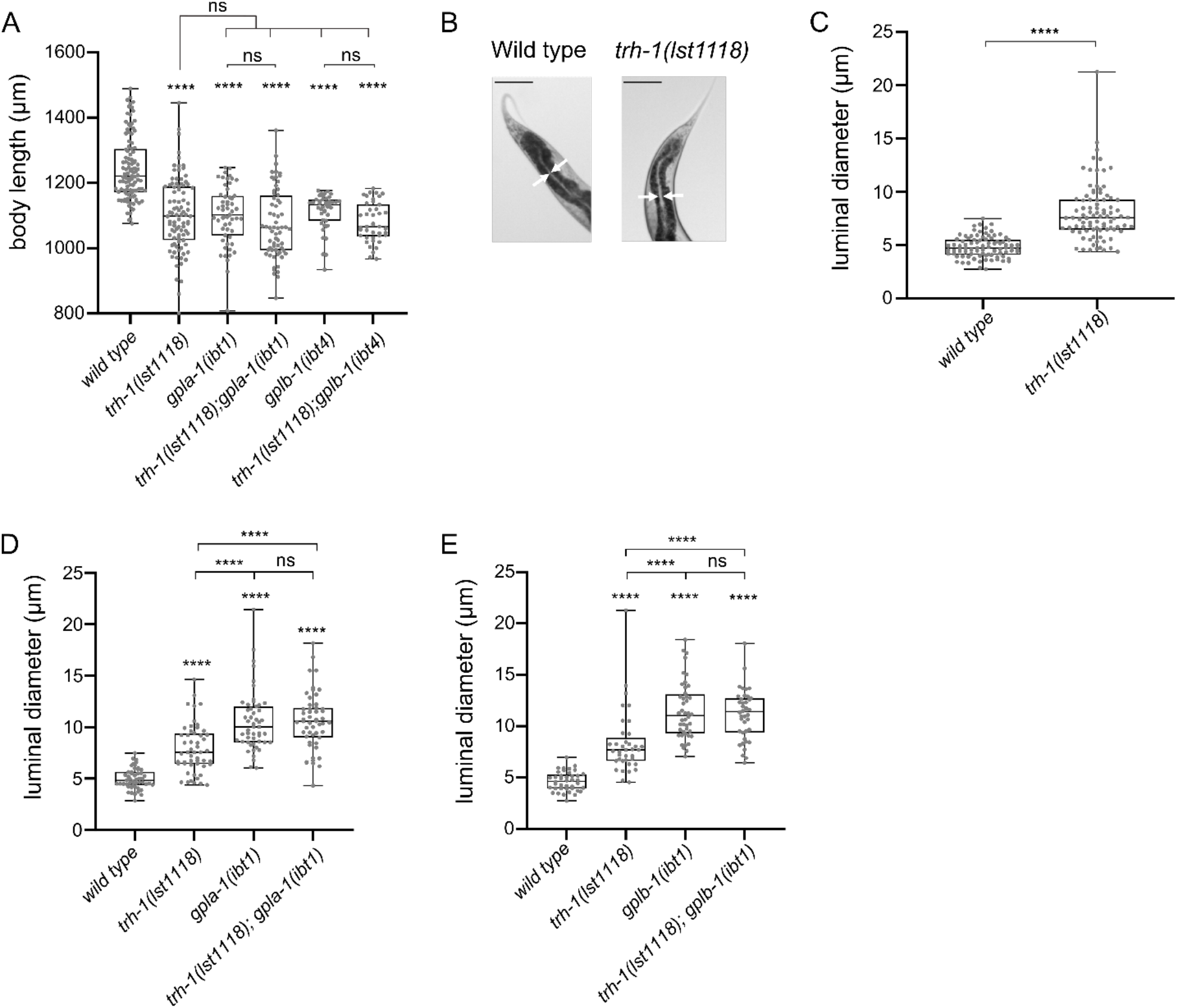
TRH-like neuropeptides and GPA2/GPB5 signaling act in the same growth-regulating pathway. (A) Double mutants defective in both the TRH-like neuropeptide precursor *trh-1* and thyrostimulin *gpla-1* or *gplb-1* do not differ in body length from the single mutants. (B-C) Mutants lacking *trh-1* show an increased luminal size of the intestine. Scale bars represent 100 μm. (D-E) Luminal width of double mutants lacking *trh-1* and *gpla-1* (D) or *gplb-1* (E) does not significantly differ from the width of single *gpla-1 or gplb-1* mutants. Boxplots indicate 25^th^ (lower boundary), 50^th^ (central line), and 75^th^ (upper boundary) percentiles. Whiskers show minimum and maximum values. For (A) and (C)-(E), each genotype was tested in at least 2 assays with 20-30 animals per trial. Data were analyzed by one-way ANOVA with Tukey’s multiple comparisons test. ****P < 0.0001; ns, not significant.

## Discussion

Phylogenetic analyses show a clear conservation of thyrostimulin GPA2 and GPB5 polypeptides across Protostomia and Deuterostomia, but their physiological roles remain elusive ^7–9,14,26,29,46^. In invertebrates, GPA2/GPB5 orthologs have only been functionally characterized in arthropods ^12,38–40^. Here, we identify a thyrostimulin-like neuroendocrine system in the nematode *C. elegans* and uncover a conserved role of this signaling pathway in the control of growth. Our findings expand our understanding of the evolutionary history and physiological actions of GPA2/GPB5 glycoprotein hormone signaling, for which we delineate a potentially ancestral role in regulating intestinal function.

The orphan receptor FSHR-1 is the sole glycoprotein hormone receptor ortholog identified in *C. elegans*, which belongs to the type A LGR family ^7,8,49^. We show that *C. elegans* GPLA-1/GPA2 and GPLB-1/GPB5 are cognate ligands of FSHR-1 *in vivo* and are able to functionally activate this receptor *in vitro*. Our results indicate that FSHR-1 has basal receptor activity, like other thyrostimulin receptors ^28,29,76– 78^. Previous work identified a sequence motif in the third intracellular loop of FSHR-1 that is likely responsible for ligand-independent signaling ^49^. In addition, we show that GPLA-1 and GPLB-1 activate FSHR-1, which stimulates cAMP signaling. These findings support the evolutionary conservation of a functional thyrostimulin-like neuroendocrine system from nematodes to humans.

In vertebrates, the beta subunits of classical glycoprotein hormones (GPB1 – 4) have a carboxytail extension, referred to as the “seatbelt”, that wraps around the alpha subunit to stabilize heterodimer configuration ^3,79,80^. However, thyrostimulin GPB5 polypeptides lack the seatbelt structure ^3,9,81^, and whether heterodimerization is required for GPA2/GPB5 signaling is debated. Moreover, differences in the expression patterns of GPA2 and GPB5, both in vertebrates and invertebrates, suggest that the two subunits may have monomeric or homodimeric functions ^21,25,27,29,38,82^. These findings have raised the question as to what the ancestral configuration of GPA2 and GPB5 hormones might have been and whether each subunit may also signal independently. *In vitro* studies of thyrostimulin receptors in humans, insects and *Amphioxus* previously found potent receptor activation only with heterodimeric or tethered GPA2/GPB5 proteins, although human GPB5 also activated TSHRs at 100-fold higher concentrations than the heterodimer ^14,23,25,28,82^. Here, we show that *C. elegans* FSHR-1 is activated by GPLA-1/GPLB-1 as well as individual GPLA-1 and GPLB-1 polypeptides. Recent analyses of the crystal structure of GPLA-1/GPLB-1 revealed that the two subunits indeed heterodimerize but are also more symmetric in structure as compared to the alpha and beta subunits of human CG and FSH ^46^. Our results are in line with this study. We find that GPLA-1 and GPLB-1 can activate FSHR-1 independently, suggesting that thyrostimulin heterodimers and other glycoprotein hormones may have evolved from an ancestral mono- or homodimeric configuration. The interaction of FSHR-1 with GPLA-1 is further corroborated by a recent study from Wang and colleagues ^83^, in which GPLA-1 was co-immunoprecipitated with the extracellular domain of FSHR-1 from transgenic *C. elegans*, suggesting that they can also bind *in vivo*. Although it remains to be understood whether heterodimerization is required for GPLA-1/GPLB-1 to exert its physiological role, we show that both subunits act in the same neuroendocrine pathway to regulate growth. In addition, GPLA-1 and GPLB-1 may exert independent functions based on our receptor activation and expression analyses. While expression of both subunits co-localizes in multiple cells, such as the enteric DVB neurons and RME motor neurons, *gpla-1* is also expressed in pharyngeal neurons and may have separate functions in the regulation of feeding behavior. GPLB-1 may additionally act as a neuroendocrine signal from non-neuronal tissues, such as the enteric muscles and the epithelial-like hmc cell.

The role of thyrostimulin GPA2/GPB5 orthologs in protostomes is largely unknown. A physiological role in growth and development, diuresis, and reproduction has previously been described in arthropods ^11–13,38,39^. In insects, the thyrostimulin receptor ortholog LGR1, like *C. elegans* FSHR-1, is abundantly expressed in the alimentary canal ^13,38^. In the adult mosquito, GPA2/GBP5 influences ion transport across the hindgut by promoting potassium secretion and inhibiting natriuresis ^13^. Thus, GPA2/GBP5 signaling may be ancestrally involved in regulating intestinal function, at least in ecdysozoans. In addition, knockdown of GPA2/GPB5 or their receptor has been shown to affect vitellogenesis in female prawns and leads to reduced reproductive success in male mosquitos ^39,41^. Thyrostimulin-like signaling may also have a role in regulating hermaphrodite reproduction in *C. elegans*, as previous research revealed that FSHR-1 is involved in germline development and fertility ^84^. In vertebrates, GPA2 and GPB5 are found in reproductive tissues as well ^10,25,82^, and a thyrostimulin-TSHR paracrine signaling system has been identified in the mammalian ovary ^32^. Besides a reproductive function, knockout of GPB5 in mice increases bone mineralization and volume, suggesting that thyrostimulin signaling is also involved in regulating mineral balance in vertebrates ^33^.

In *C. elegans*, we found that thyrostimulin-like signaling promotes growth via activation of FSHR-1 in the intestine and glial cells. We show that GPLA-1 and GPLB-1 are expressed in enteric neurons associated with the *C. elegans* pharynx and gut, and identify the intestine as a main target site for GPA2/GPB5 signaling, as loss of GPLA-1/GPLB-1 or FSHR-1 causes intestinal bloating. Distention of the intestinal lumen has been observed in *C. elegans* upon pathogenic infection and in mutants defective in the defecation motor program ^62,85,86^. Consistent with these studies, we find that the defecation cycle period is significantly longer in *gpla-1, gplb-1* and *fshr-1* mutants than in wild type animals. The defecation motor program has important functions in the uptake of dietary nutrients. For example, defecation defects have been linked with aberrant fat metabolism and a reduced uptake of fatty acids from the intestinal lumen ^87^. Such nutritional deficits may impair growth. Indeed, we find that other mutants defective in the defecation motor program, like thyrostimulin mutants, have a shorter body length.

How thyrostimulin-like signaling might regulate the periodicity of the defecation cycle remains to be determined. Previous research revealed that the period is set by spontaneous calcium oscillations in the intestinal cells. These trigger pH oscillations and neuropeptide release in the gut, which mainly control the execution of the defecation motor program ^62,88,89^. Mutations that disrupt intestinal calcium or pH oscillations cause defects in period length and variability ^56,88,90^. In addition, cycle length is influenced by various environmental and internal factors, such as temperature ^91^, food ^54^, reactive oxygen species ^57^, and infection ^92^. We speculate that GPLA-1/GPLB-1 signaling, through activation of FSHR-1, might regulate rhythmic calcium or pH signaling in intestinal cells. This is supported by a previous study in which loss of GPLA-1 and the putative lipid-binding protein GHI-1 was shown to partially suppress defecation defects in mutants lacking *flr-1*, an acid-sensing member of the Degenerin/Epithelial Sodium Channel (DEG/ENaC) superfamily that is required for intestinal calcium and pH oscillations ^15,93^. Interestingly, changes in intestinal pH also affect pathogen susceptibility, and mutants of both *fshr-1* and *gpla-1* were previously found to be more susceptible to infection ^15,60,94^. FSHR-1 has been shown to protect *C. elegans* from pathogenic bacteria by inducing pathogen avoidance, upregulating stress response genes, and delaying intestinal accumulation of ingested pathogens ^59,60,85,95^, although it remains unknown whether these functions are mediated by GPLA-1/GPLB-1 signaling. In addition, FSHR-1 has been implicated in host responses to oxidative and freezing-thawing stress ^60,83^. Whether FSHR-1 regulates the defecation motor program upon infection or oxidative stress has not yet been determined, but it is conceivable that thyrostimulin-like signaling may also regulate protective immune and stress responses in *C. elegans*.

In mammals, the hypothalamic neuropeptide TRH is a releasing factor for growth-promoting hormones, like GH and TSH from the pituitary gland ^74^. Vertebrate TSH and thyrostimulin both induce the release of growth-stimulating thyroid hormones ^2,10,25^. Evidence for the existence of these hormones in nematodes is lacking ^96^. However, we find that GPLA-1/GPLB-1 and TRH-related neuropeptides, encoded by the *trh-1* gene, act in the same genetic pathway to regulate body size in *C. elegans*. Mutations in *trh-1* also cause bloating of the intestine, similar to the phenotypes observed in mutants defective in GPLA-1/GPLB-1 signaling. TRH-like neuropeptides and their receptors are widely conserved across protostomian and deuterostomian animals ^43,97–99^. It will be interesting to see whether the TRH system influences the release of invertebrate glycoprotein hormones, similar to its function in vertebrates, or whether release of GPA2/GPB5 is mediated by different factors.

Taken together, our findings lend further support to the ancient origin of the thyrostimulin GPA2/GPB5 glycoprotein hormone system that may have evolved from mono- or homodimeric antecedents predating the split of the vertebrate lineage. Our genetic analysis of the GPA2/GPB5 system in *C. elegans* suggests that this neuroendocrine axis together with TRH-like neuropeptides constitute a conserved pathway that regulates growth. Our results also provide a molecular basis for further investigations into a potentially ancestral role of GPA2/GPB5 signaling in regulating intestinal function.

## Acknowledgements

We thank the *Caenorhabditis* Genetics Center (CGC, University of Minnesota), funded by the NIH Office of Research Infrastructure Programs (P40 OD010440), for providing strains; all members of the Beets lab for experimental advice; N. De Fruyt for help on statistics; M. Christiaens and A. Kieswetter for technical assistance; and J. Kowalski, W.R Schafer, E. Kaulich and D. Walker for advice and reagents. This research was supported by the Research Foundation Flanders (FWO) grant G0C0618N and KU Leuven Research Council grant C16/19/003 (to I.B. and L.S.). S.V.D. is a fellow of the Research Foundation Flanders (FWO) and J.W. is supported by a postdoctoral fellowship of the KU Leuven Research Council.

## Author contributions

S.K., M.N.I, J.W., L.S. and I.B. designed the research. S.K., M.N.I., S.V.D. and E.V. performed the experiments. S.K. and M.N.I. analyzed data. I.B. and L.S. supervised the project and were responsible for funding acquisition. S.K. and I.B. wrote the paper. All authors revised and edited the manuscript.

## Materials and Methods

### *C. elegans* strains and maintenance

All *C. elegans* strains were maintained at 20°C on nematode growth medium (NGM) plates seeded with *Escherichia coli* OP50 bacteria. All experiments were performed using 1-day adult hermaphrodites, unless mentioned otherwise. Wild-type (N2-Bristol), IBE7 *flr-2 (ut5)*, IBE24 *flr-2 (ok2127)*, IBE1 *fshr-1 (ok778)* and EG1285 *lin-15B&lin-15A (n765) oxIs12 [unc-47p::GFP + lin-15(+)]* strains were obtained from the *Caenorhabditis* Genetics Center (CGC, University of Minnesota). CB156 *unc-25 (e156)* and EG276 *exp-1 (ox276)* strains were a kind gift from the lab of W.R. Schafer (Laboratory of Molecular Biology, Cambridge, UK). Deletion alleles for *gpla-1* (*ibt1*) and *gplb-1* (*ibt4*) were obtained by CRISPR/Cas9 genome editing, as described below. A full list of strains used in this study can be found in Table S1.

### Molecular biology

All fluorescent reporter and rescue constructs were made using the MultiSite Gateway Three-Fragment cloning system (Invitrogen). Genomic DNA or cDNA sequences of *fshr-1, gpla-1*, and *gplb-1* were cloned in front of a SL2 trans-splicing site and fluorescent reporter gene.

Putative promoter sequences for *fshr-1* (3110 bp), *gpla-1* (4058 bp), and *gplb-1* (3589 bp) were cloned from genomic DNA of wild-type *C. elegans*. Tissue-specific rescue transgenes were generated using promoter regions of *rab-3* (1208 bp) ^65,66^, *rgef-1* (3660 bp) ^69^, *ges-1* (3313 bp, kind gift from the lab of W.R. Schafer, Cambridge, UK) ^61–63^, and *mir-228* (2217 bp) ^64^. All constructs have a *let-858* 3’UTR except the *rgef-1* rescue construct, which was made using a *tbb-2* 3’UTR.

The plasmid for heterologous expression of *fshr-1* in HEK cells was obtained by directionally cloning the *fshr-1* cDNA into the pcDNA3.1/V5-His-TOPO vector (Invitrogen). The cDNA sequence of the *fshr-1a* gene isoform was amplified by PCR using cDNA from mix-staged wild-type *C. elegans* as template.

The forward primer included a ‘CACC’ sequence at the 5’ end that introduced a partial Kozak sequence for increased translation efficiency in mammalian cells.

### CRISPR/Cas9 knockout strain generation

The *gpla-1(ibt1)* and *gplb-1(ibt4)* knockout alleles were generated by CRISPR/Cas9-mediated deletion of the *gpla-1* or *gplb-1* open reading frames. For each gene, two crRNA sequences were designed by looking for PAM sites (NGG) that were in the vicinity of the double stranded break site. After analysis of putative off-target sites (http://crispor.tefor.net/), we selected the highest scoring sequences based on their predicted on-target activities. Repair templates were designed to be in-frame and were codon-optimized for *C. elegans* (https://www.genscript.com/tools/codon-frequency-table). To simplify selection of the successfully edited worms, we used a Co-CRISPR technique in which the *dpy-10* gene was also knocked out by CRISPR editing ^100^. We used the CRISPR/Cas9 protocol as described by Mello *et al*. ^100^ for the preparation of injection mixes containing all required components (Table S2). After injection of young adult hermaphrodites, F1 progeny that showed the rolling and/or dumpy phenotype were transferred to individual NGM plates and checked for gene deletion by PCR and subsequent gel-electrophoresis. Successful deletion of the genes was confirmed by sequencing.

The *gpla-1 (ibt3* [His::GPLA-1]) knockin allele was made by CRISPR/Cas9 mediated insertion of a His-tag at the N-terminus of the *gpla-1* open reading frame, using a similar strategy as described above and the crRNA and repair template listed in Table S2.

Below, the sequences of the gene knockouts and knockins are shown with flanks on the (-) strand. Exons are in grey, with start and stop codons in red, the remaining gene sequences after CRISPR gene editing are underlined, and with the inserted gene sequence in blue.

*gpla-1(ibt1)* deletion:

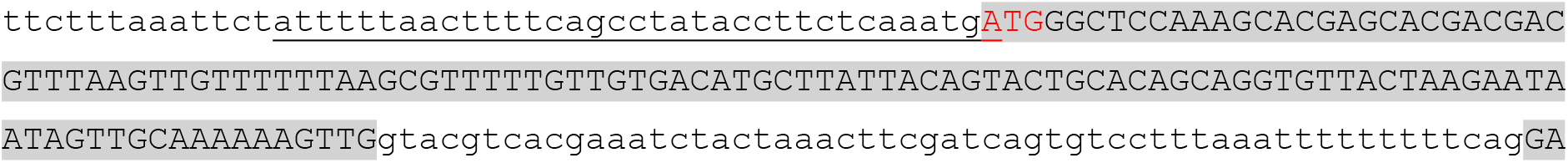
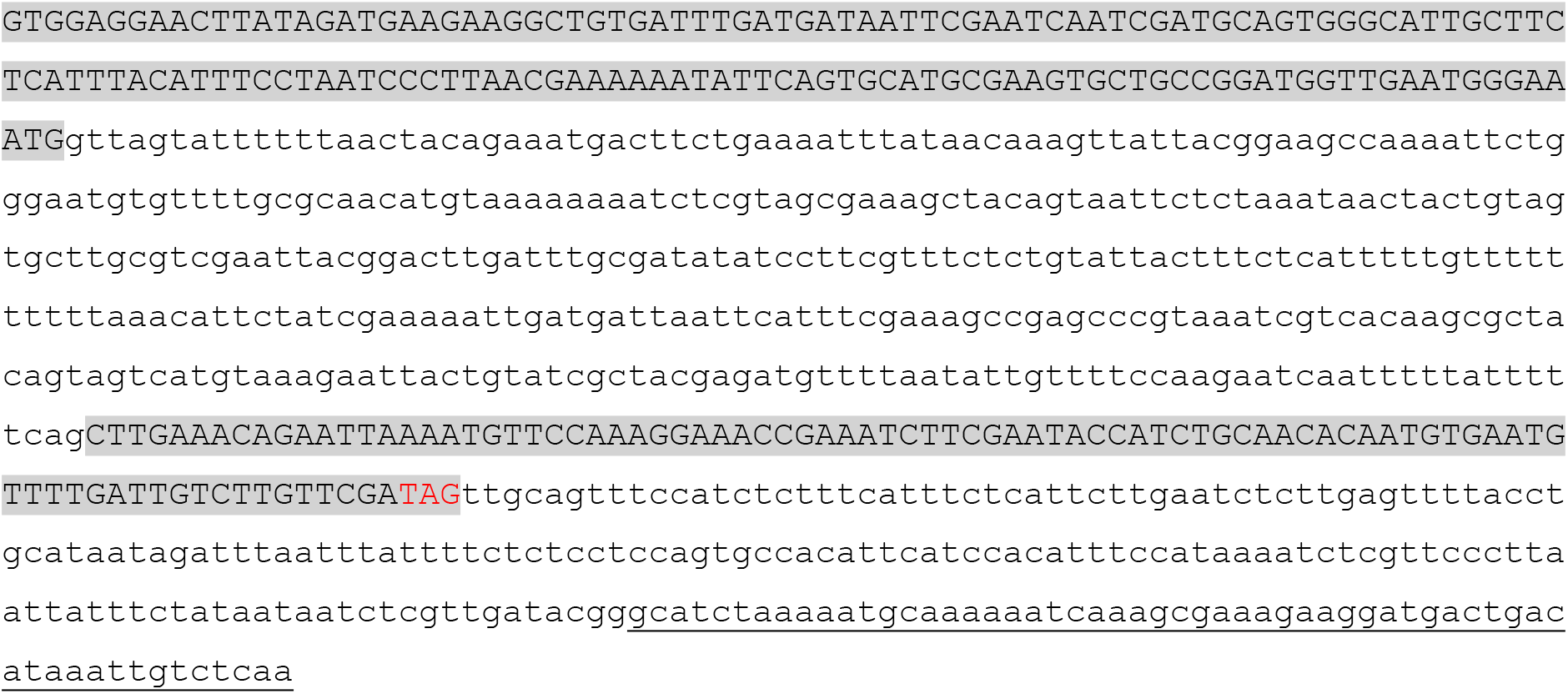

*gplb-1(ibt4)* deletion:

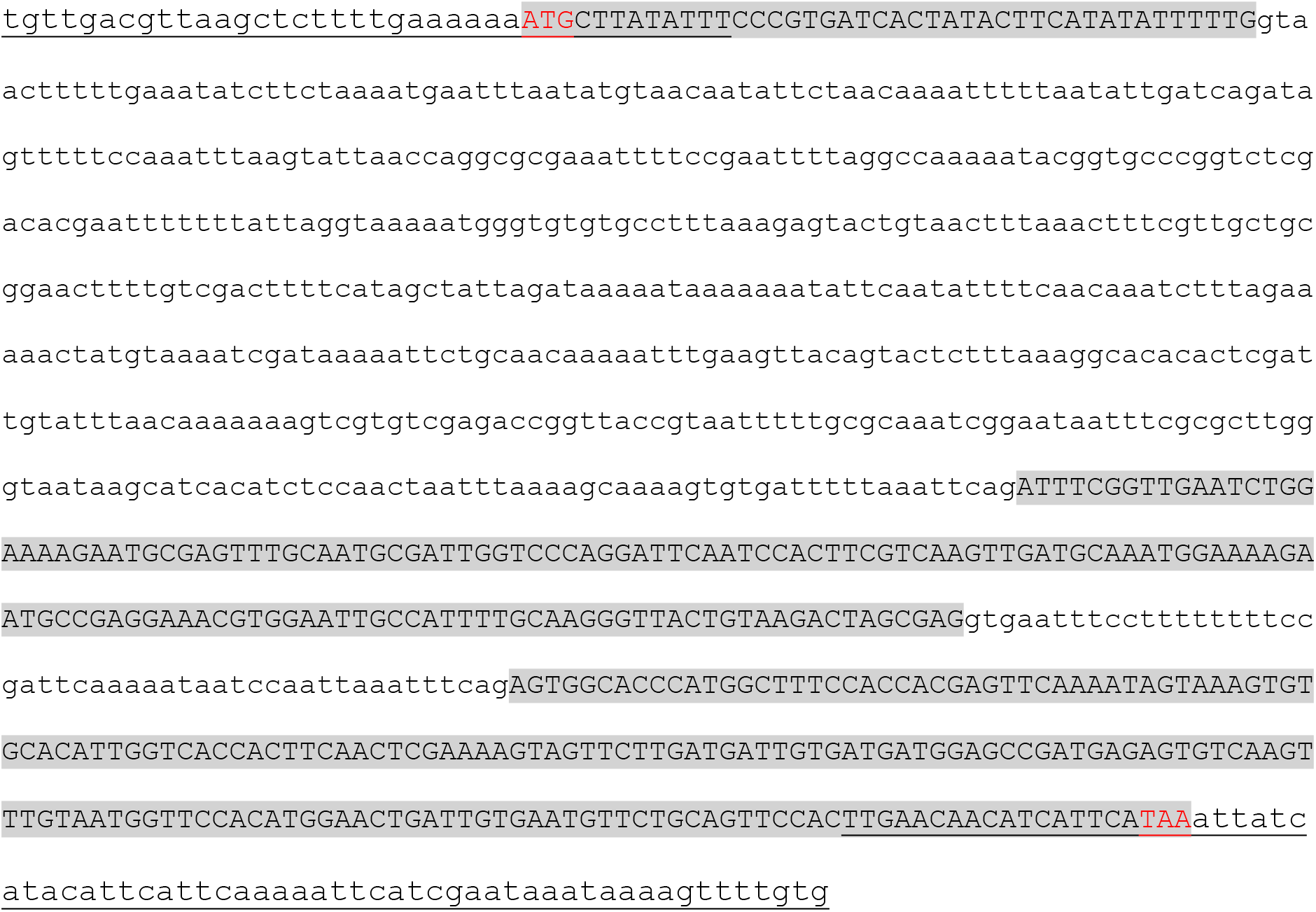

*gpla-1(ibt3* [His::GPLA-1]) insertion:

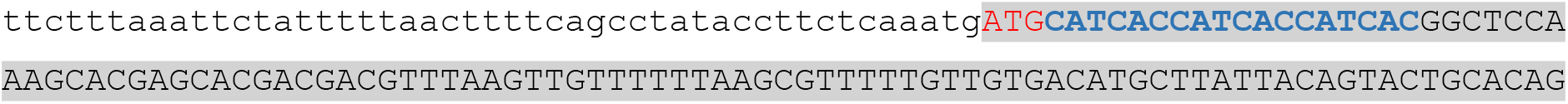
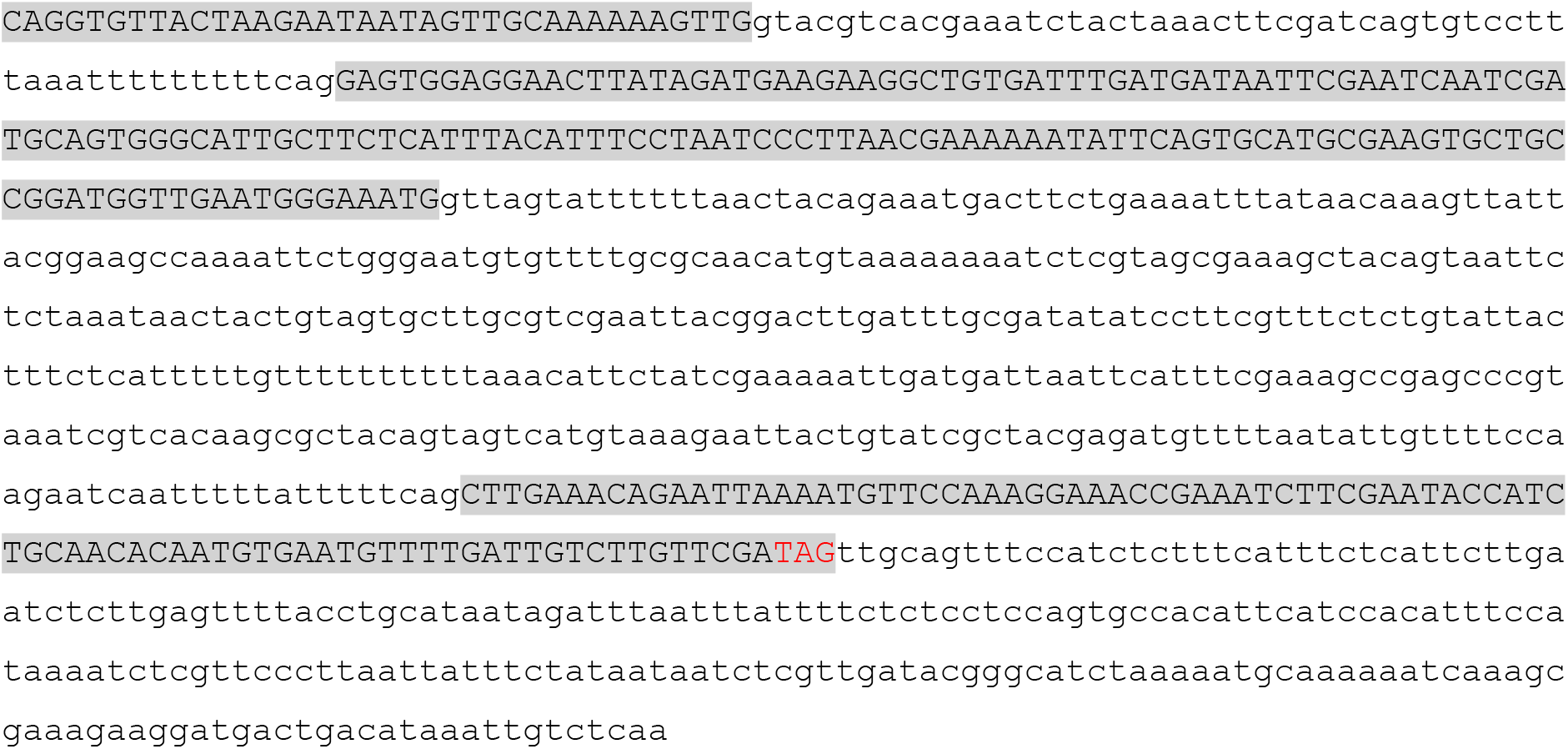

## Transgenesis and expression pattern analysis

Germline transformations were carried out by injecting constructs into the syncytial gonad of young adult worms together with a co-injection marker (*myo-2p::mCherry, unc-122p::dsRed*, or *unc-122p::GFP*) and 1-kb DNA ladder (Thermo Scientific) as carrier DNA.

Expression patterns of fluorescent reporter transgenes were visualized by an Olympus confocal microscope (FluoView FV1000, IX81) and resulting Z-stack projections were created and analyzed with ImarisViewer (v9.7.2) software. Adult worms were immobilized by mounting in 50 mM Sodium Azide solution in M9 buffer on a 2% agarose pad and covered with a glass cover slip. Expression patterns were confirmed in at least two independent transgenic strains. DVB and RME expression was confirmed by crossing with marker strain EG1285, which marks GABAergic neurons ^101^. Other cells expressing *gpla-1, gplb-1*, or *fshr-1* were identified based on their position and morphology.

## Body size and luminal width quantification

Body size parameters (length, width, volume) of day one adults were measured using the custom WormSizer MATLAB script (https://github.com/jwatteyne/WormSizer). Synchronized L1 larvae were placed onto freshly seeded NGM plates one day post-synchronization and kept at 20°C until assaying.

After 65 hours on food, 20 to 30 well-fed day one adults were transferred from the plates and anesthetized in 20 µL of 10 mM tetramisole hydrochloride solution (Sigma-Aldrich) in Milli-Q water on a 2% agarose pad. Images were captured with a ZEISS Axio Observer.Z1 at 5x magnification. Body size was measured from images of anesthetized day 1 adult worms on at least two independent days. After skeletonization, delineating the worm’s outline and midline, the worm’s total length and middle width were calculated using the calibration factor (pixels/µm) of the pictures. For the volume estimation, the midline of the worm was separated in 30 segments of the same size. The volume for each segment was determined by using the formula for the frustum of a cone, taking the natural shape of a worm into consideration. The summation of all these volumes was described as the worm’s total volume. To assess growth throughout adulthood, body size measurements were executed for 5 subsequent days (day 1 to day 5 adulthood) for at least two independent time periods.

Intestinal lumen diameter was quantified from images taken for body size measurements of day 1 adult worms. Using the ImageJ software (version 1.53s), luminal width was determined by measuring the average width (in µm) of the intestine at two different points in the posterior lumen. Body size parameters and luminal widths were plotted, and significance levels were calculated with GraphPad Prism 8 software.

### Recombinant protein synthesis

Recombinant proteins were synthesized and purified by OriGene™ Technologies (Rockville, USA). To generate recombinant GPLA-1/GPLB-1, HEK293 cells were transfected with expression vectors encoding *C. elegans* N-terminal 6xHis-tagged GPLA-1 and non-tagged GPLB-1. Recombinant proteins were purified from cell lysate by affinity chromatography in PBS buffer (PBS and 10% glycerol, pH 7.3) and analyzed by Tricine-SDS-PAGE on a 8 – 20% gel in Tris-Glycine running buffer and by subsequent Western blot using anti-His monoclonal antibodies. A similar approach was used to synthesize individual GPLA-1 and GPLB-1 proteins. N-terminal 6xHis-tagged GPLA-1 was expressed in HEK293 cells and purified by affinity chromatography. Non-tagged GPLB-1 was expressed in HEK cells and purified using ion exchange chromatography. Both recombinant GPLA-1 and GPLB-1 protein samples were analyzed by Tricine-SDS-PAGE.

### cAMP bioluminescence assay

The cAMP bioluminescence assay quantifies ligand-induced bioluminescent responses by measuring changes in intracellular cAMP levels after receptor activation. HEK293T cells were cultured in monolayer at 37°C in a humidified atmosphere with 5% CO2, in Dulbecco’s Modified Eagle’s Medium (DMEM)/Nutrient F-12 Ham (Gibco) supplemented with 10% heat inactivated Fetal Bovine Serum (FBS, Sigma-Aldrich) and 1% Penicillin/Streptomycin (Gibco). Co-transfection with the cAMP indicator CRE(6x)-luciferase and pcDNA3.1/*fshr-1a* was done in a 1:1 ratio when cells reached 60-70% confluency. Transfection medium contained Opti-MEM (Gibco, Life Technologies), CRE(6x)-luciferase and receptor plasmid DNA, Plus Reagent (Invitrogen) and Lipofectamine LTX (Invitrogen). One day post-transfection, fresh culture medium was added to the transfected cells. Two days post transfection, each well of a Bio-One CELLSTAR™ 96-well, Flat Bottom Microplate (Greiner) was loaded with a dilution series of recombinant protein compounds in 200 µM 3-isobutyl-1-methylxanthine (IBMX), which inhibits cAMP hydrolysis. Compound buffer (PBS with 10% glycerol) without hormone was added to the IBMX medium as a ligand-free negative control. Cells were detached, counted, pelleted and resuspended to a concentration of 10^6^ cells/mL in IBMX medium. Each well of the compound plate was supplemented with 50 µL of the cell suspension (50 000 cells/well) and the plate was incubated at 37°C for 3.5 h. Cells were then loaded with 100 µL SteadyLitePlus substrate (PerkinElmer) and incubated on a shaking plate for 15 minutes under dark conditions at RT. Finally, luminescence was measured twice for 5s (at 0s and 5s) per well at 469 nm on a Mithras LB 940 luminometer (Berthold Technologies). Measurements were performed in triplicate in at least 4 independent experiments. Luminescence values were plotted and significance levels were calculated with GraphPad Prism 8 software.

### Defecation motor program (DMP) assay

The defecation motor program was analyzed as previously described ^102^. Animals were synchronized by letting day one adults lay eggs for 2-3 hours, before removing them from the NGM plates, and allowing the eggs to develop into day one adults. One day prior to the DMP assay, L4 animals were picked onto a freshly seeded NGM plate and all strains were blinded. After 15-17 hours, worms were acclimated on the microscope stage for ∼10 minutes. At least 10 consecutive defecation cycles were observed from each animal on a Nikon stereomicroscope. Cycle length, aBoc and expulsion events were scored, and defecation behaviour was logged using BORIS (Behavioral Observation Research Interactive Software). Animals were assayed alongside wild type and at least one defecation-defective control. The custom DefecationAnalysis R script was applied to measure the defecation parameters of day one adults (https://github.com/NathanDeFruyt/DefecationAnalysisBeetsLab). The length of one DMP cycle was defined by the time elapsed between two subsequent contractions of the posterior body wall muscles (pBocs). The average cycle length was calculated over 10 defecation cycles for each worm. aBoc frequency was defined as the ratio of aBoc over pBoc. Defecation parameters were plotted and significance levels were calculated with GraphPad Prism 8 software.

### Sequence alignment of GPA2 and GPB5 orthologs

The *H. Sapiens* GPA2 and GPB5 and *C. elegans* GPLA-1 and GPLB-1 sequences were used as queries to identify sequences with highest similarity in selected model systems representative for their phylum by the Basic Local Alignment Search Tool (BLAST) of the National Center for Biotechnology Information (NCBI) database. The multiple sequence alignment program Clustal Omega (https://www.ebi.ac.uk/Tools/msa/clustalo/) was used to align sequences, and conserved residues were identified using BoxShade (https://embnet.vital-it.ch/software/BOX_form.html).

**Figure S1.**
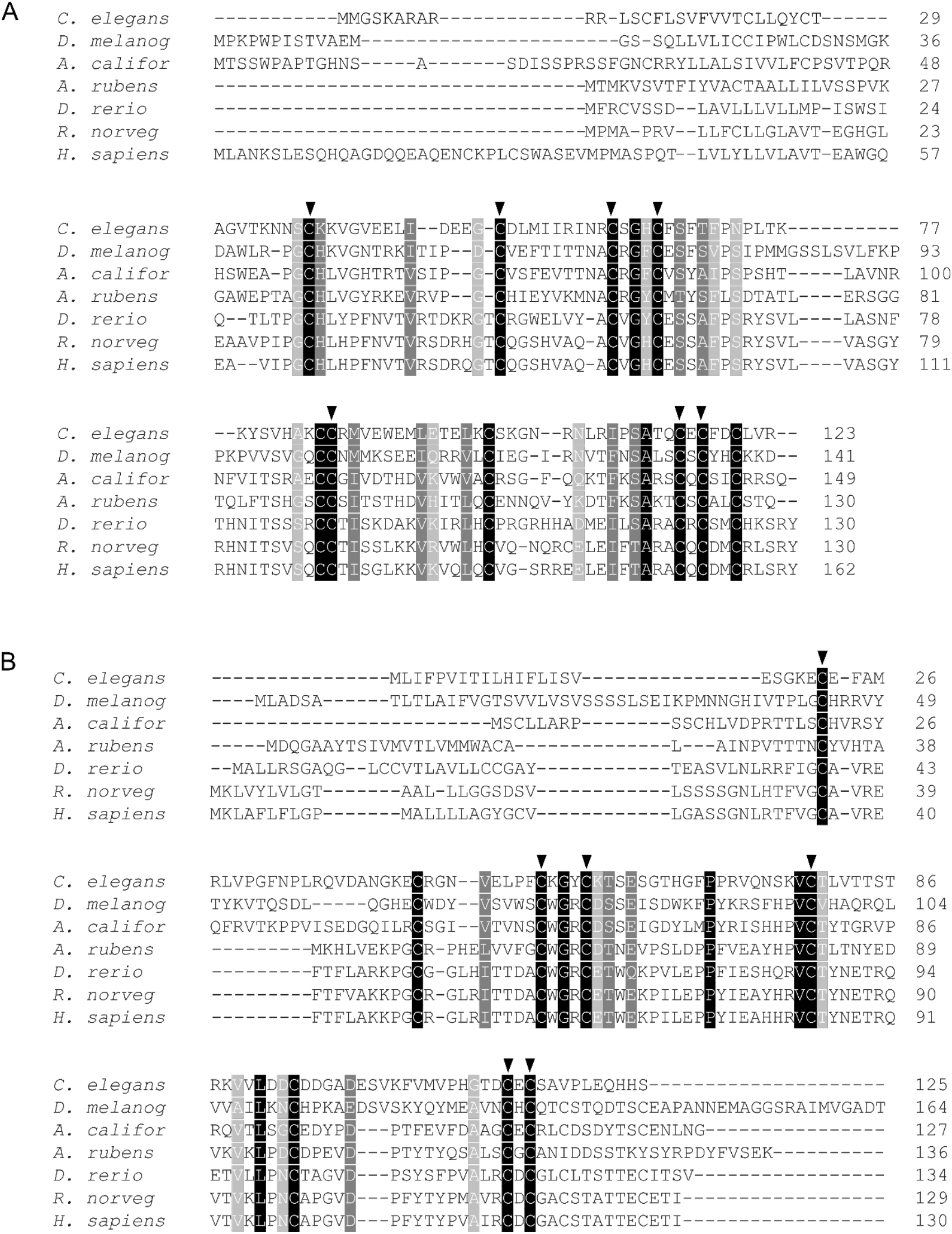
*C. elegans* GPLA-1 and GPLB-1 show sequence similarity with thyrostimulin GPA2 and GPB5. Protein sequence alignment of *C. elegans* GPLA-1 (A) and GPLB-1 (B) with GPA2 and GPB5 orthologs, respectively, in representative protostomian and deuterostomian species, including *Homo sapiens, Rattus norvegicus, Danio rerio, Asterias rubens, Aplysia californica*, and *Drosophila melanogaster* (Table S3). Six cysteine residues forming the typical cysteine knot structure of glycoprotein hormone subunits are highly conserved and indicated with black arrow heads. Black shading depicts full amino acid conservation, dark grey indicates groups of amino acids with strong similar properties, light grey depicts weakly similar amino acids.

**Figure S2.**
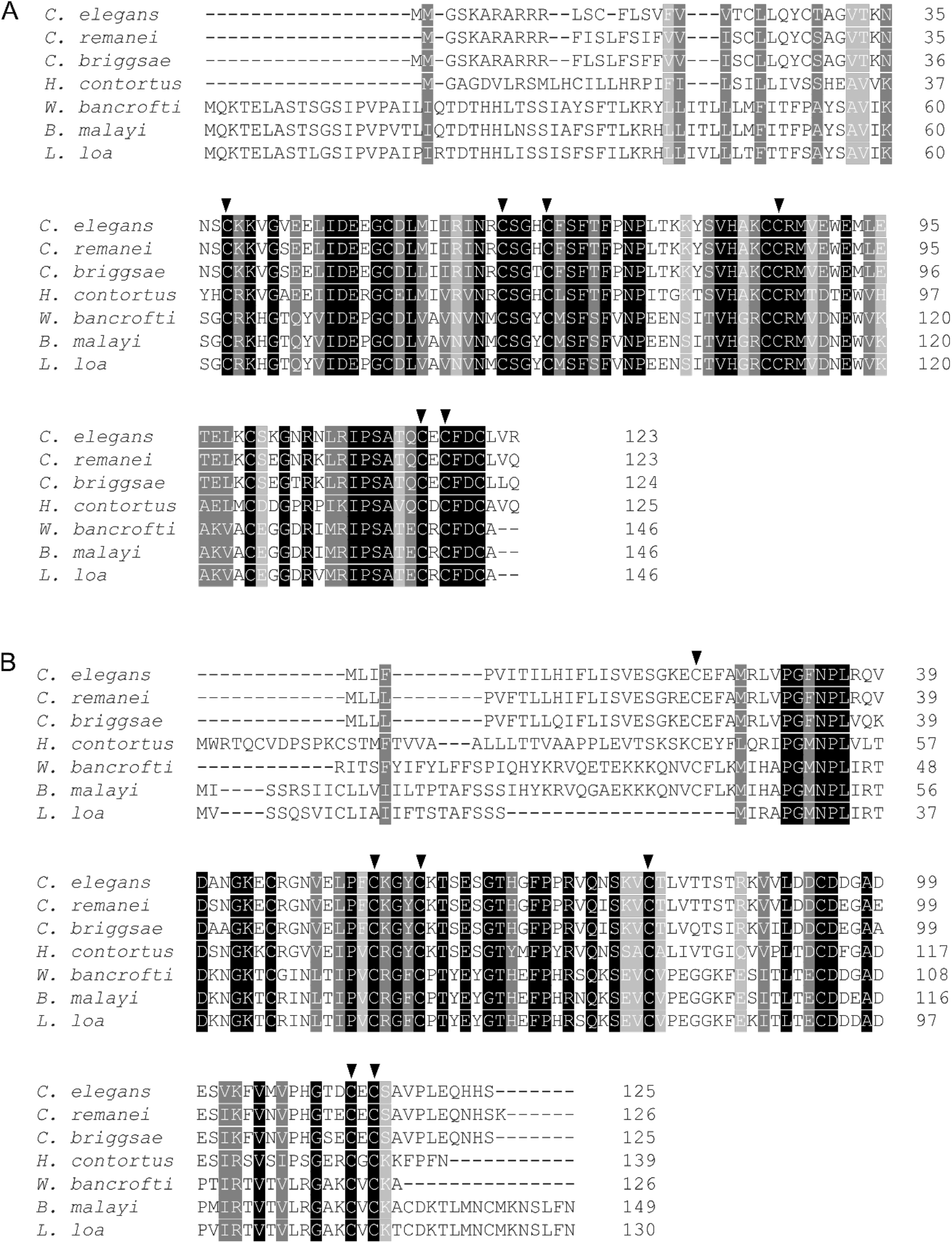
Conservation of thyrostimulin GPA2 and GPB5 subunits in nematodes. Protein sequence alignment of GPLA-1/GPA2 (A) and GPLB-1/GPB5 (B) sequences in nematode species, including *Caenorhabditis elegans, Caenorhabditis remanei, Caenorhabditis briggsae, Haemonchus contortus, Wuchereria bancrofti, Brugia malayi*, and *Loa loa* (Table S3). Symbols and shading as in figure S1.

**Figure S3.**
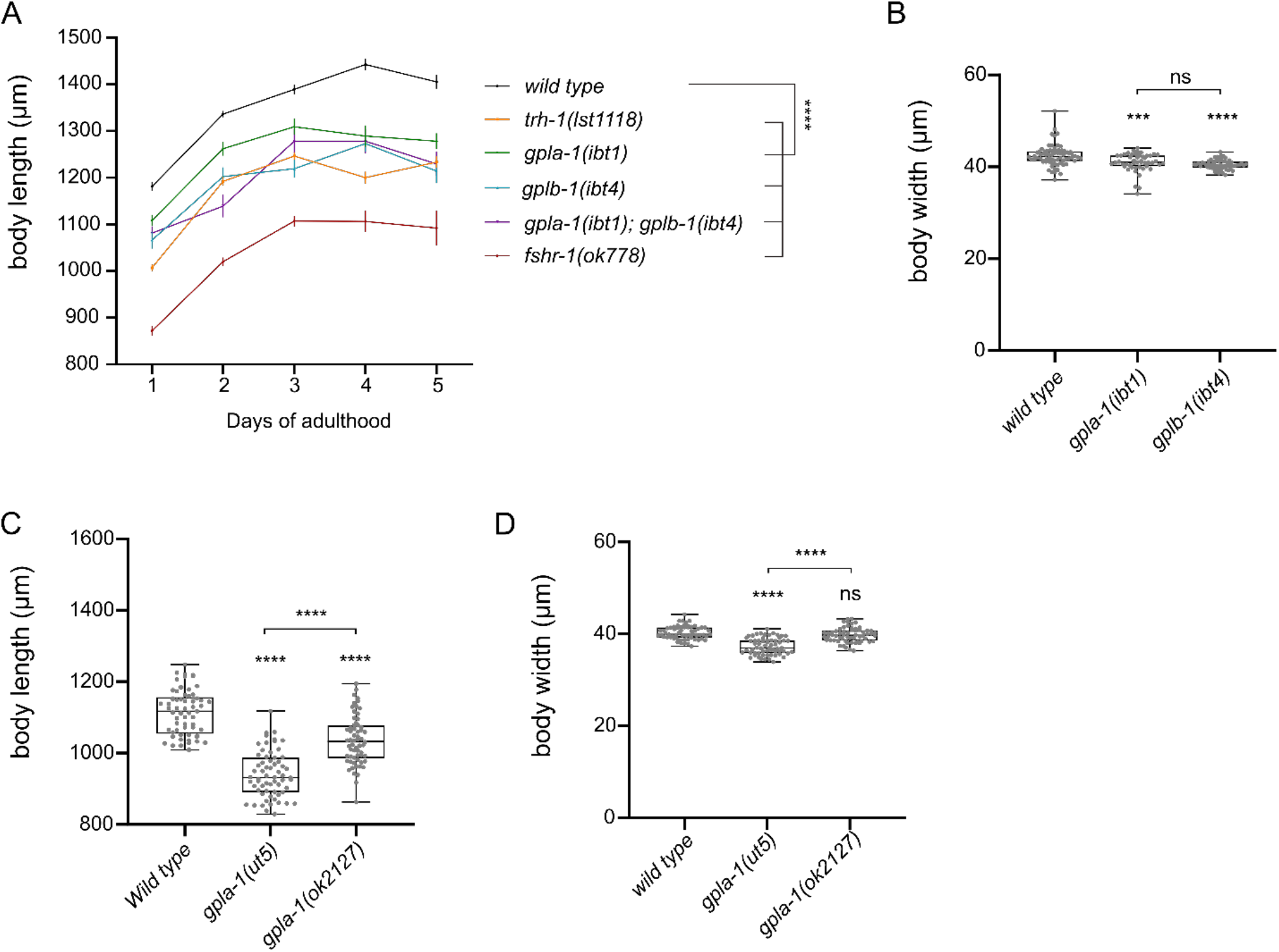
Mutants lacking thyrostimulin-like signaling show body size defects throughout adulthood. (A) Null mutants of *gpla-1, gplb-1* and *fshr-1* have a significantly shorter body length than wild-type animals throughout adult life, from day 1 to day 5 of adulthood. Mutants of the TRH-like neuropeptide *precursor trh-1* also have a shorter body length43 and were used as positive control. Data is plotted as the mean body length ± SEM. For each genotype, around 20 animals were tested on each day of adulthood in at least 2 assays. (B) Comparison of body width of *gpla-1* and *gplb-1* mutants. (C-D) Comparison of body length (C) and width (D) of different *gpla-1* mutant alleles. Boxplots indicate 25^th^ (lower boundary), 50^th^ (central line), and 75^th^ (upper boundary) percentiles. Whiskers show minimum and maximum values. For (B)-(D), each genotype was tested in 3 assays with 20-30 animals per trial. Data were analyzed by one-way ANOVA with Tukey’s multiple comparisons test. ***P < 0.001; ****P < 0.0001; ns, not significant.

**Figure S4.**
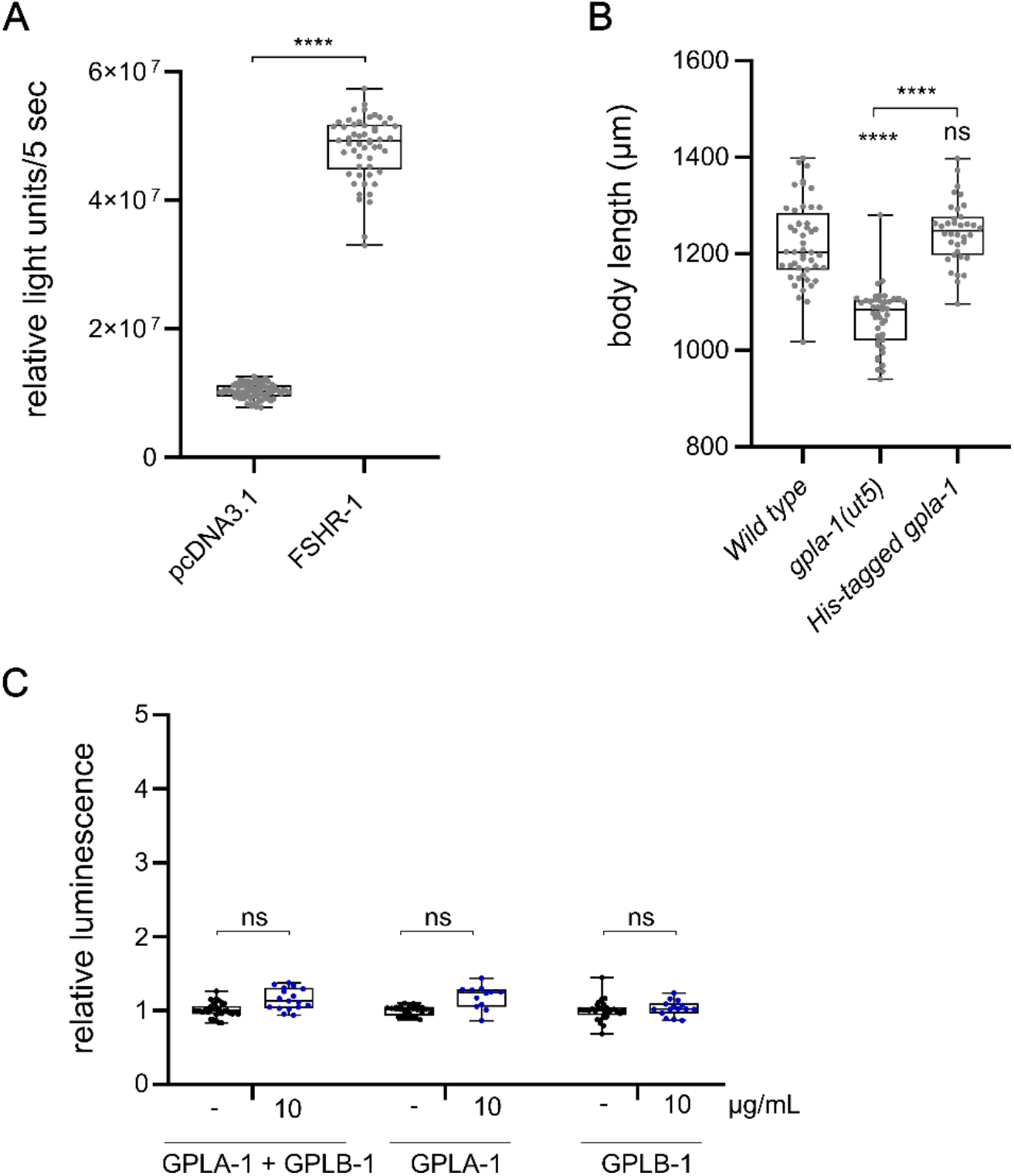
FSHR-1 shows basal activity in a cAMP-based receptor activation assay *in vitro*. (A) HEK cells expressing FSHR-1 show significantly higher cAMP-induced luminescence than cells transfected with an empty pcDNA3.1 vector control. Unpaired t test, n = 3 assays. (B) Transgenic *C. elegans* expressing N-terminal His-tagged GPLA-1 do not differ from wild-type animals in body length. Boxplots indicate 25^th^ (lower boundary), 50^th^ (central line), and 75^th^ (upper boundary) percentiles. Whiskers show minimum and maximum values. Each genotype was tested in 2 assays with 20-30 animals per trial. One-way ANOVA with Tukey’s multiple comparisons test. (C) Recombinant GPLA-1 and GPLB-1 proteins do not induce a significant increase in cAMP-mediated luminescence in cells transfected with an empty pcDNA3.1 vector control. Data are shown as relative luminescence after normalization to the ligand-free control. Two-way ANOVA with Dunnett’s multiple comparisons test, ligand free controls, n ≥ 4 assays. ****P < 0.0001; ns, not significant.

**Figure S5.**
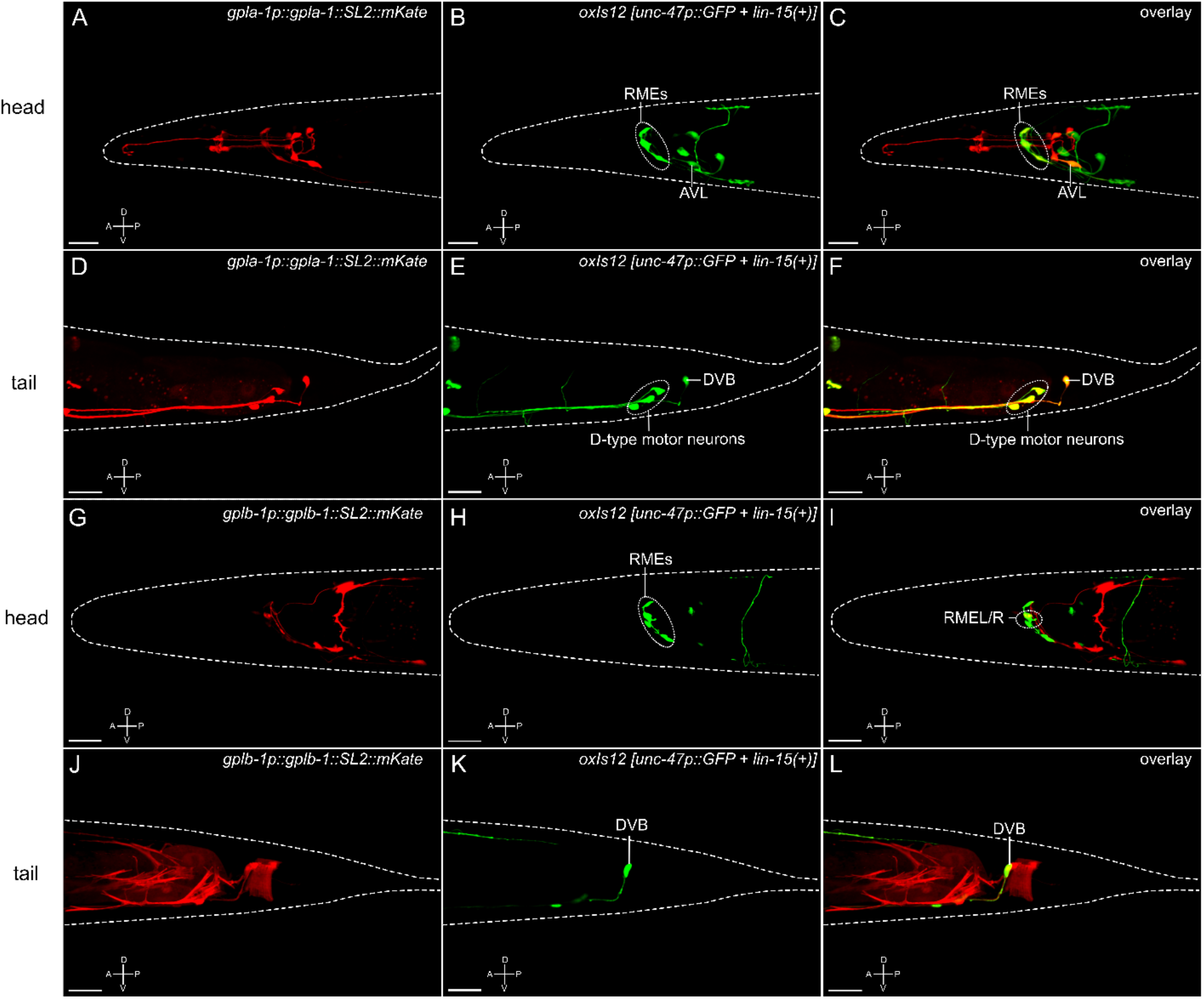
Identification of *gpla-1* and *gplb-1* expression sites in *C. elegans*. Labeled confocal Z-stack projections of *gpla-1* and *gplb-1* fluorescent reporter transgenes. Overlap with the GABAergic reporter strain EG1285 (*oxIs12 [unc-47p::GFP + lin-15(+)]*), marking GABAergic neurons, shows co-localization in RME and DVB neurons for *gpla-1* and *gplb-1*, and additionally in AVL and D-type motor neurons for *gpla-1*. A; anterior, P; posterior, D; dorsal, V; ventral orientation. Scale bars represent 20 µm.

**Figure S6.**
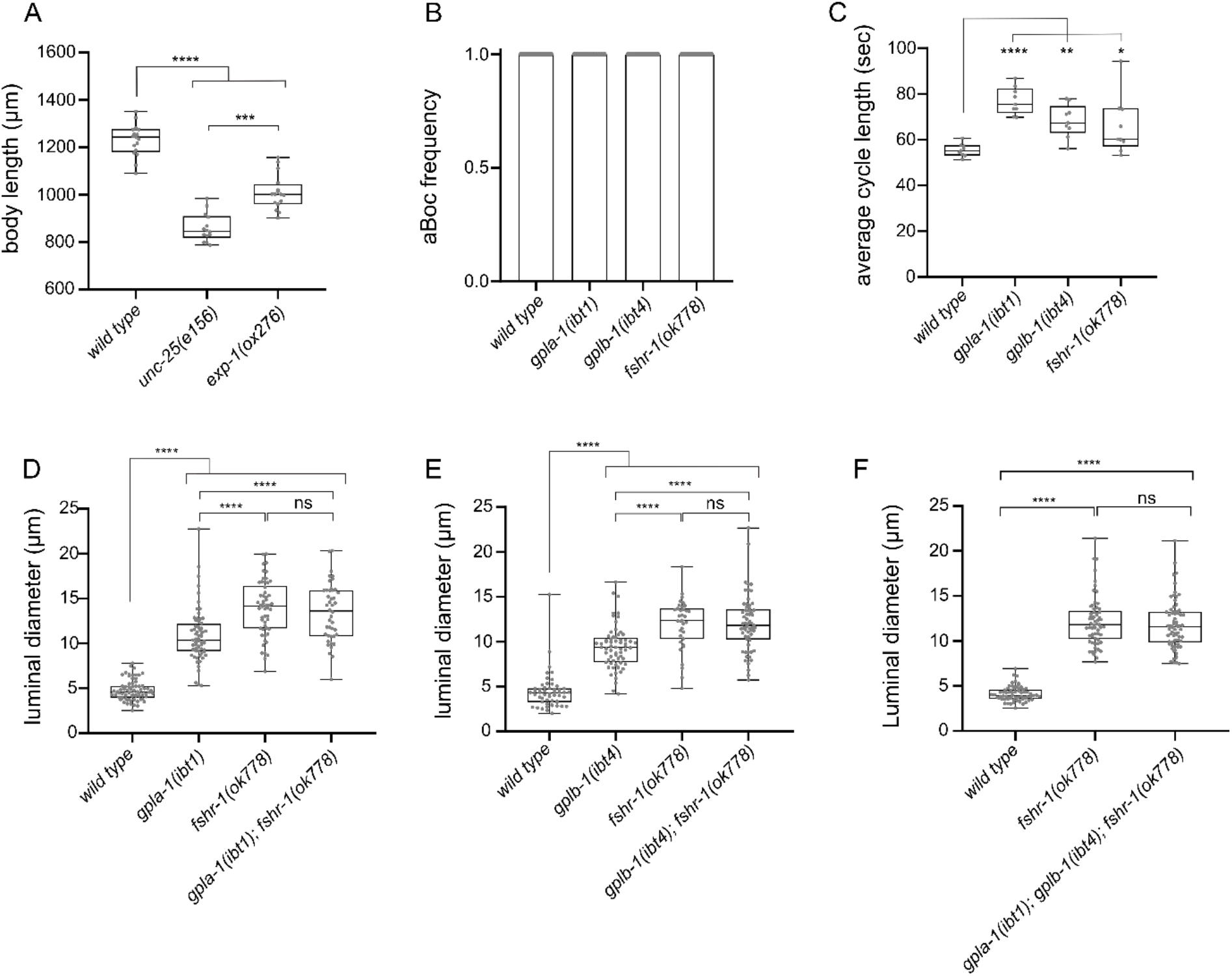
Mutants impaired in thyrostimulin-like signaling show defects in the defecation motor program and intestinal lumen size. (A) GABAergic transmission mutants *unc-25(e156)* and *exp-1(ox276)* that are defective in the defecation motor program70–72 also show defects in body length. n ≥ 15 animals per strain (B) The aBoc frequency *of gpla-1, gplb-1*, and *fshr-1* mutants does not differ from wild-type animals. Bar graphs depict the mean aBoc frequency ± SEM. One-way ANOVA with Dunnett’s multiple comparisons test. (C) *gpla-1, gplb-1*, and *fshr-1* mutants show a significantly increased defecation cycle length compared to wild type. One-way ANOVA with Dunnett’s multiple comparisons test. (D-E) Luminal width of double mutants lacking *fshr-1* and *gpla-1* (D) or *gplb-1* (E) does not significantly differ from the luminal width of single *fshr-1* mutants. One-way ANOVA with Tukey’s multiple comparisons test. (F) The intestinal lumen of mutants lacking *gpla-1, gplb-1* and *fshr-1* is not significantly wider than that of single *fshr-1* mutants. One-way ANOVA with Tukey’s multiple comparisons test. For (A) and (C)-(F), boxplots indicate 25^th^ (lower boundary), 50^th^ (central line), and 75^th^ (upper boundary) percentiles. Whiskers show minimum and maximum values. For (B)-(C), each genotype was tested in 3 assays with 3 animals per trial. For (D)-(F), each genotype was tested in 2 assays with 20-30 animals per trial. *P < 0.05; **P < 0.01; ***P < 0.001; ****P < 0.0001; ns, not significant.

**Table S1:**
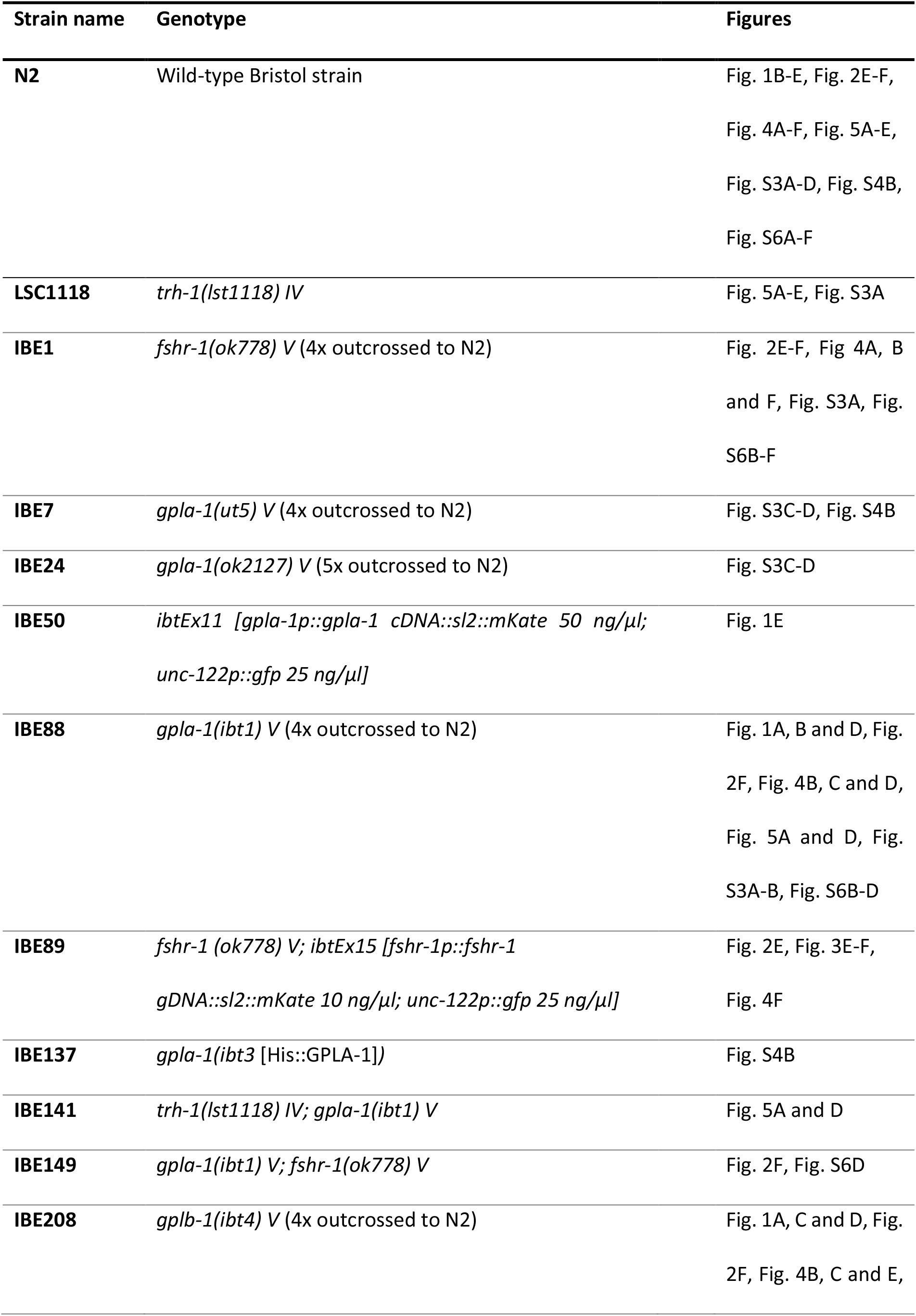
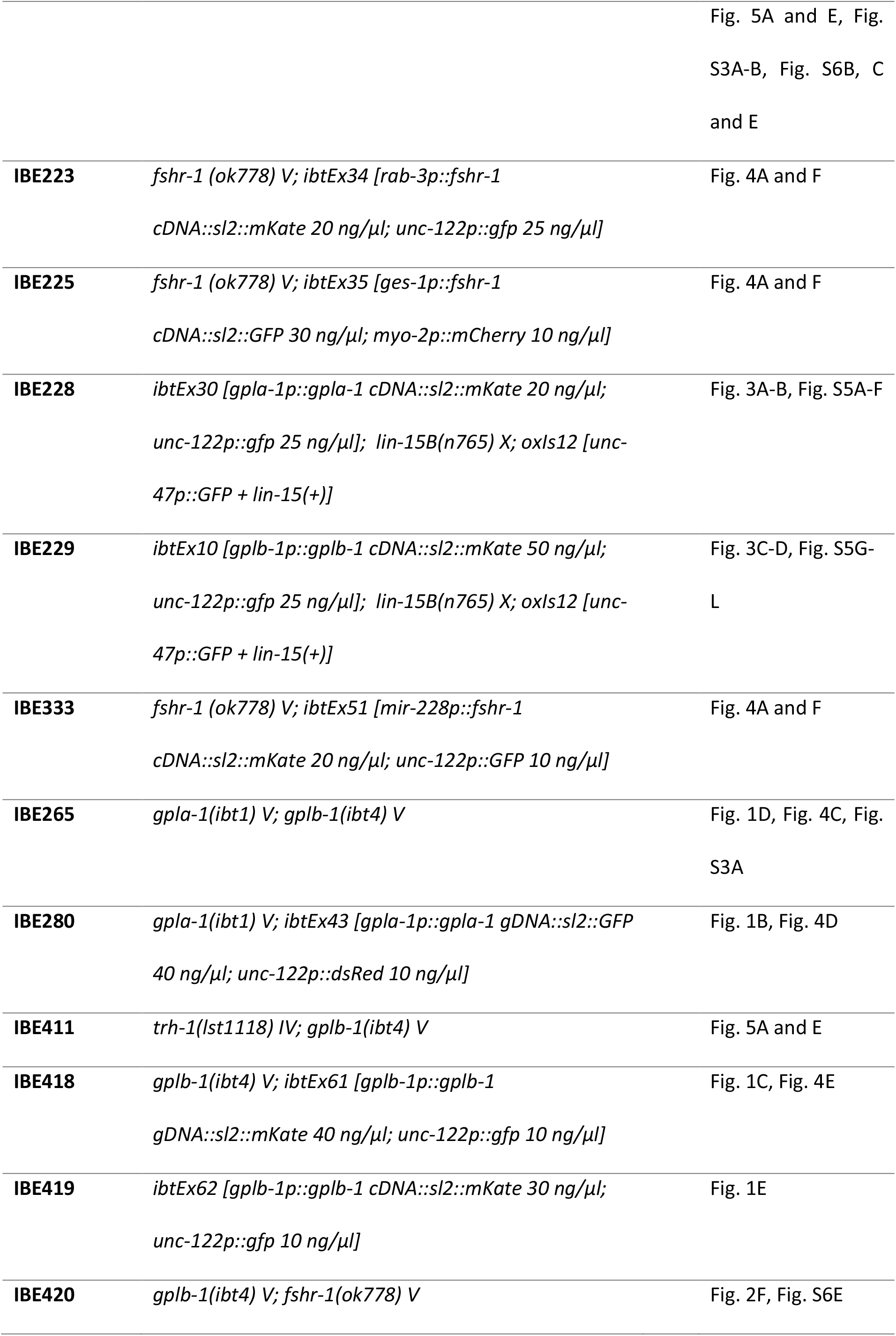
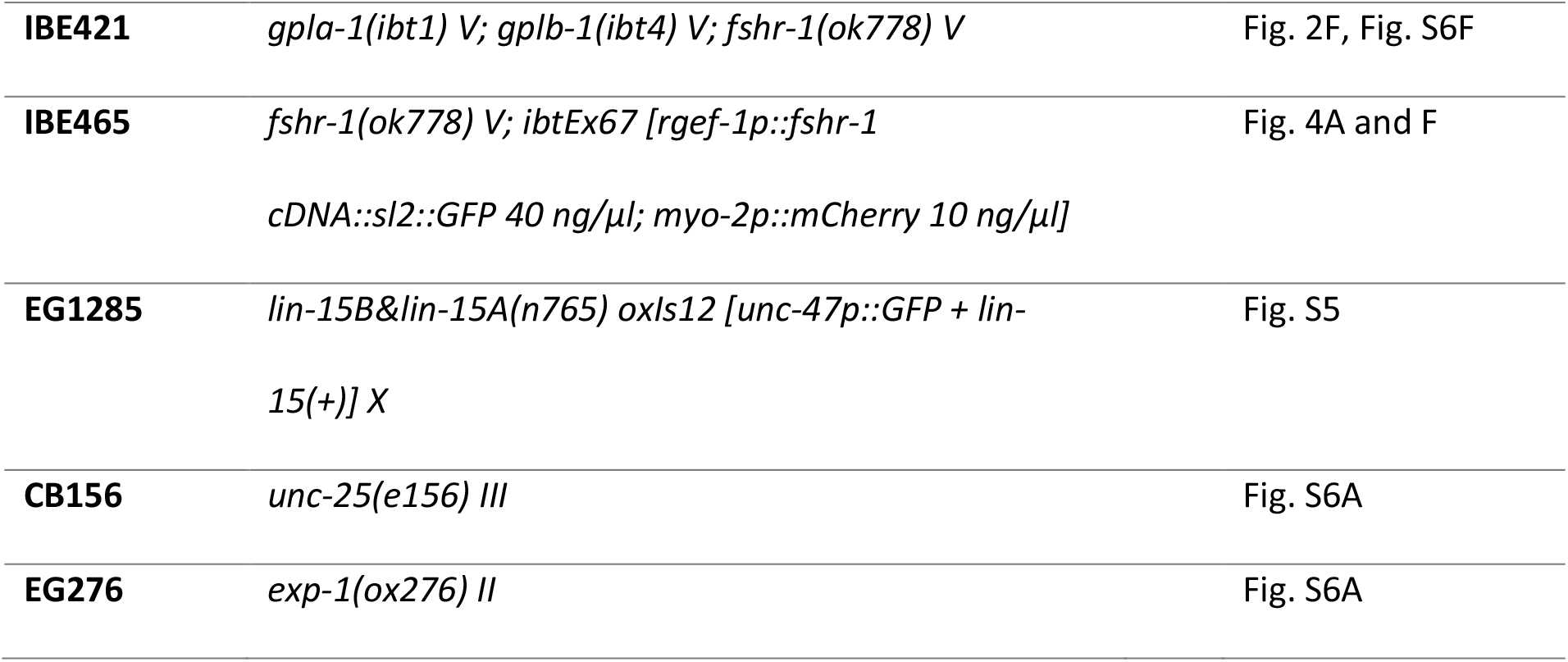
Overview of *C. elegans* strains used in this study.

**Table S2:**
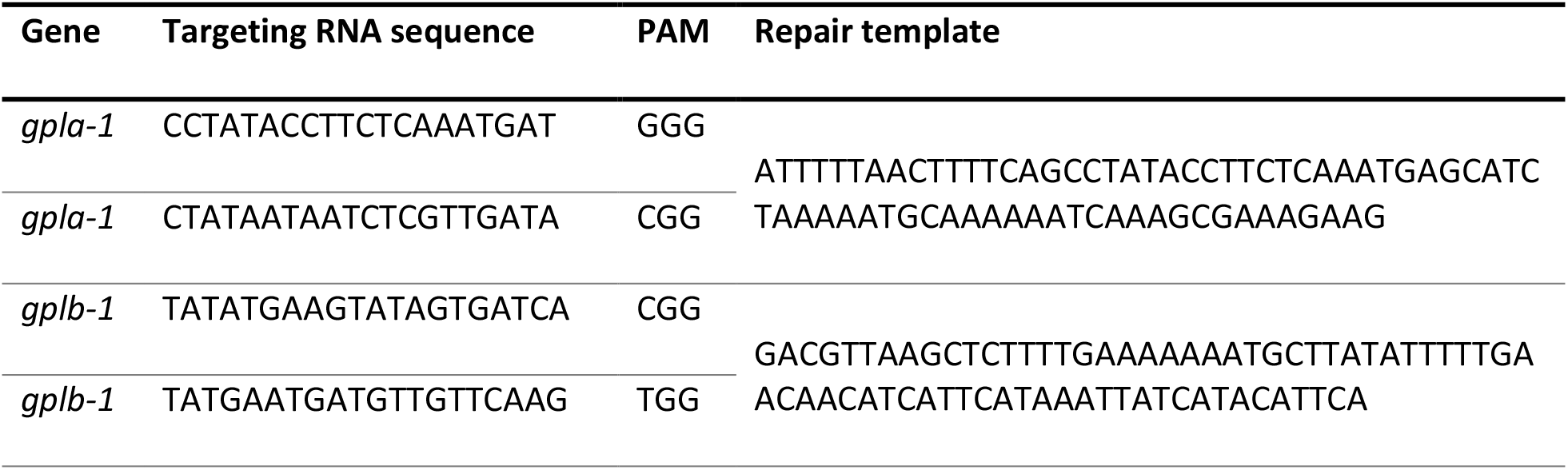
CRISPR RNA sequences used to target *gpla-1* and *gplb-1* with the respective 3ʹ PAM sequences and repair sequences.

**Table S3:**
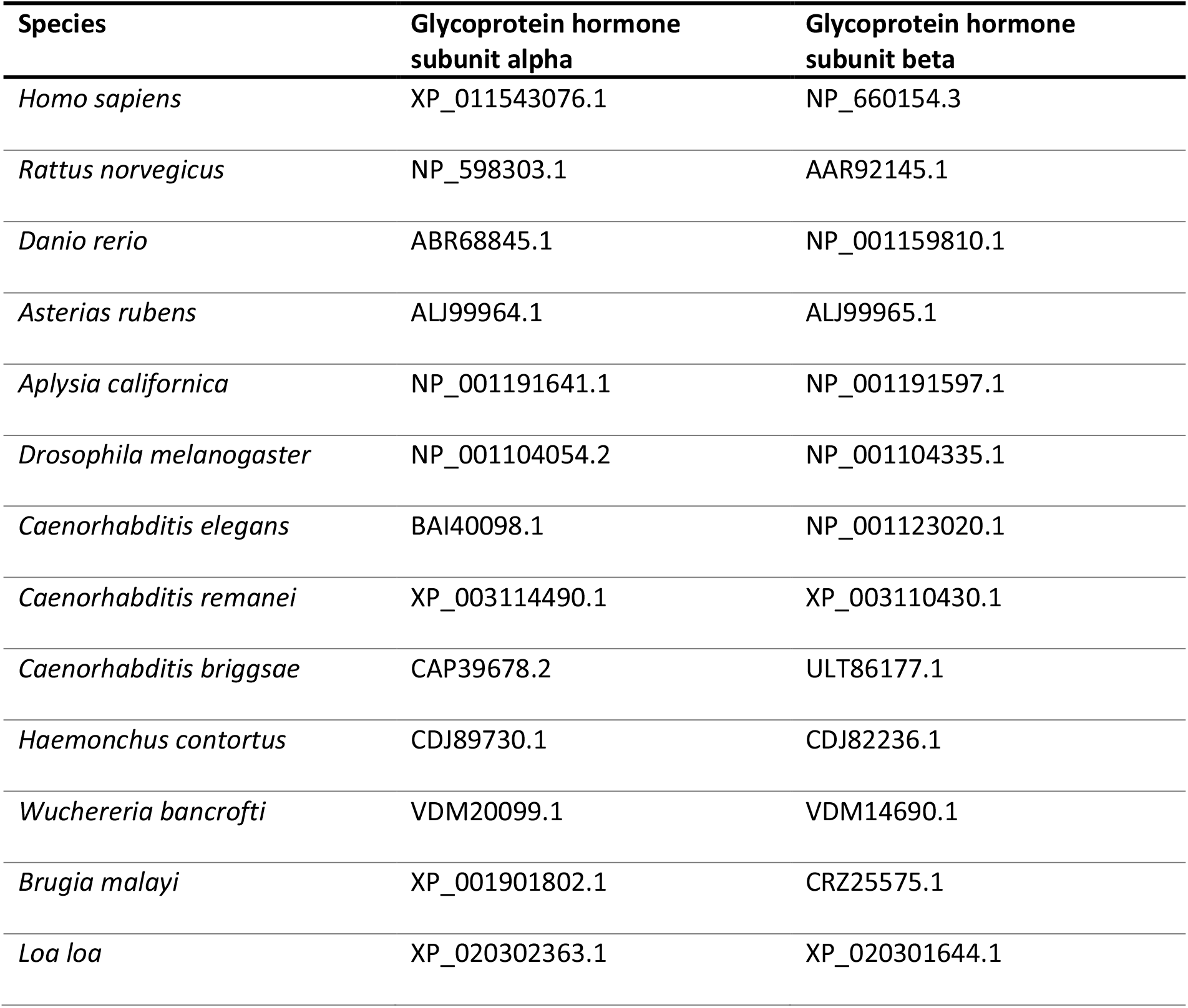
Accession numbers of the alpha and beta glycoprotein hormone subunits used in this study.

| Species | Glycoprotein hormone subunit alpha | Glycoprotein hormone subunit beta |
| --- | --- | --- |
| *Homo sapiens* | XP_011543076.1 | NP_660154.3 |
| *Rattus norvegicus* | NP_598303.1 | AAR92145.1 |
| *Danio rerio* | ABR68845.1 | NP_001159810.1 |
| *Asterias rubens* | ALJ99964.1 | ALJ99965.1 |
| *Aplysia californica* | NP_001191641.1 | NP_001191597.1 |
| *Drosophila melanogaster* | NP_001104054.2 | NP_001104335.1 |
| *Caenorhabditis elegans* | BAI40098.1 | NP_001123020.1 |
| *Caenorhabditis remanei* | XP_003114490.1 | XP_003110430.1 |
| *Caenorhabditis briggsae* | CAP39678.2 | ULT86177.1 |
| *Haemonchus contortus* | CDJ89730.1 | CDJ82236.1 |
| *Wuchereria bancrofti* | VDM20099.1 | VDM14690.1 |
| *Brugia malayi* | XP_001901802.1 | CRZ25575.1 |
| *Loa loa* | XP_020302363.1 | XP_020301644.1 |

